# Multiple layers of innate immune response antagonism of SARS-CoV-2

**DOI:** 10.1101/2024.01.29.577695

**Authors:** Fuchun Zhou, Sivakumar Periasamy, Nathaniel D. Jackson, Wan Sze Cheng, Ruben Soto Acosta, Philipp A. Ilinykh, Chengjin Ye, Shailendra Chauhan, German Nudelman, Elena Zaslavsky, Steven G. Widen, Luis Martinez-Sobrido, Stuart C. Sealfon, Alexander Bukreyev

## Abstract

Several SARS-CoV-2 proteins have been shown to counteract the host innate immune response, mostly using *in vitro* protein expression, which may not fully reflect their role in the context of viral infection. In addition, while each viral protein was characterized in a different experimental system, their relative contribution in immunosuppression remains unclear. Here we used a SARS-CoV-2 bacterial artificial chromosome with *en passant* mutagenesis to recover a panel of twelve infectious recombinant SARS-CoV-2 viruses, each with mutations in either NSP1, NSP2, NSP3, NSP6, NSP12, NSP13, NSP14, NSP15, NSP16, ORF3a, ORF6 or ORF8. We used the interferon-stimulated response element (ISRE)-driven luciferase assay in 293T-ACE2/TMPRSS2 cells to test the panel, demonstrating that mutations in many proteins, especially in NSP1 and NSP15, increased the type I interferon response relative to the parental wild-type virus. RNA-seq analysis of mutant-virus infected Calu-3 cells showed that the mutations in NSP1 or NSP15 lead to higher expression of multiple genes involved in innate immune response, cytokine-mediated signaling and regulation of lymphocyte proliferation. Furthermore, mutations in either NSP1 or NSP15 resulted in a greater maturation of human monocyte-derived dendritic cells *in vitro*. Infection of K18 hACE2 transgenic mice with either NSP1 or NSP15 mutated viruses demonstrated attentuated respiratory tract replication. Analysis of lung immune cells from infected mice by single-cell RNA-seq identified 15 populations of major myeloid and lymphoid cells with changes in the pattern of their activation associated with viral infection. The effects of mutations in NSP1 or NSP15 on these responses are consistent with differences in the immunosuppressive mechanisms utilized by the two proteins. Overall, these data demonstrate different and redundant mechanisms of innate immune antagonism by SARS-CoV-2 and suppression of activation of antigen presenting cells and T and B lymphocytes mediated by multiple viral proteins.

**AUTHOR SUMMARY:** The mechanisms by which SARS-CoV-2 and its proteins modulate host immunity, specifically the interferon response, are still not clear. We generated twelve infectious SARS-CoV-2 viruses with mutations in individual proteins and demonstrated that many of them have interferon-antagonizing activity and immunosuppressive effects in human cells and in the K18 hACE mouse model of infection. We idemtified distinct and redundant mechanisms of immunosuppression of SARS-CoV-2 mediated by multiple individual viral proteins, with 9 out of the 12 tested proteins showing some immunosuppressive effect in at least one experimental system. The demonstrated immunosuppressive effects extend from the innate response to immune cells to pathologic changes *in vivo*. Importantly, this work shows, for the first time, a comparison of the effects of multiple viral proteins in the context of authentic viral infection, rather than in a surrogate system, and shows the relative contribution of each viral protein under identical experimental conditions. Overall, our data indicates that SARS-CoV-2 antagonizes multiple immune mechanisms, particularly type I interferon signaling, activation of innate immune cells and T and B lymphocyte functions with the greatest effects due to NSP1 and NSP15.

## INTRODUCTION

The COVID-19 pandemic, caused by SARS-CoV-2, has caused more than 772 million confirmed cases of human disease including almost 7 million deaths (December 2023) (1). Since the first report on occurrence and rise of COVID-19 cases, SARS-CoV-2 has shown a powerful transmission potential that has contributed to the deadly pandemic. This virus has continuously evolved into variants of concern (VOC) with a higher ability for transmission and continuing the pandemic beyond two years (2). SARS-CoV-2, similarly to SARS-CoV, enters mammalian cells using the angiotensin-converting enzyme 2 (ACE2) receptor, which is abundant in the respiratory and intestinal epithelial cells (3). The virus causes diffuse alveolar damage and COVID-19 associated acute respiratory distress syndrome (ARDS) through dysfunctional immune responses (4). In addition, it also causes acute tissue injury, particularly in the liver and kidneys, coagulopathies (disseminated intravascular coagulation and fibrin thrombi formation), thrombocytopathy and pulmonary embolism (4). The large RNA genome of SARS-CoV-2, consisting of 14 open reading frames (ORF), encodes 4 structural proteins (S, E, M and N), 16 non-structural proteins (NSP1-16), and eleven accessory proteins (ORF3a, ORF3b, ORF3c, ORF3d, ORF6, ORF7a, ORF7b, ORF8, ORF9b, ORF9c and ORF10). The 16 NSPs are from processing of the polyprotein precursors by the viral NSP3 and NSP5, which have the activity of papain-like protease (PLpro) and the 3C-like protease (3CLpro) respectively. NSP3 is responsible for the proteolytic cleavage of NSP1-4, and NSP5 for the processing of other cleavage sites that results in NSP5-16 (5). While NSPs play a critical role in viral replication, the accessory proteins do not, but both groups of proteins play an important role in viral pathogenesis and modulating the host immune response (6).

Interferons (IFN) are important secreted host proteins with strong anti-viral functions. Type I IFNs (IFN-I), which includes IFNα and IFNβ are produced by many cells in response to viral infections including SARS-CoV-2 (7). Data from clinical patients indicate that low induction of IFN-I at local and systemic level correlates with the severity of COVID-19 (8–10). In some patients with severe COVID-19, high nasal viral titers correlate with low IFN-I levels, and in some cases with high titers of autoantibodies against IFNs (11). This suggests that an appropriate induction of high titers of IFNs enables epithelial cells to inhibit SARS-CoV-2 replication and growth at nasopharyngeal and other mucosal sites (11, 12). However, SARS-CoV-2 antagonizes the innate host defense mechanisms including the inhibition of IFN-I production and signaling (13) allowing the virus to replicate exponentially causing severe tissue pathologies.

The mechanisms by which SARS-CoV-2 and its proteins modulate host immunity, specifically the IFN response, are still not clear. Recent studies support the formulation that NSPs and accessory proteins of SARS-CoV-2 antagonize IFN-I signaling (14–26). The role of SARS-CoV-2 proteins in IFN-I antagonism has been investigated by transfection of cells with plasmids overexpressing viral proteins and luciferase reporter assay. Lei et al. found that NSP1, NSP3, NSP12, NSP13, NSP14, ORF3a, and ORF6 inhibit induction of IFN-β by Sendai virus infection, whereas NSP2 and S protein induced IFN-β (23). Yuen et al. found that NSP13, NSP14, NSP15 and ORF6 suppress IFN-I production and signaling (25). Using a similar system, Shemesh et al. found that NSP1, NSP5, NSP6, NSP15, ORF6 and ORF7b block MAVS-induced (but not TRIF-induced) IFNβ production (18). Specifically, NSP6, ORF6, and ORF7b blocked MAVS-induced IFNβ-promoter activity leading to reduction of IFNβ mRNA and protein expression. SARS-CoV-2 ORF6 and ORF7b gene products also blocked IFNβ production via MAVS-dependent pathway involving both MDA5 and RIG-I signaling (18). Investigation of SARS-CoV-2 and host protein interactions using affinity purification and mass spectrophotometry (AP-MS) found that many viral proteins including NSP13, NSP15, ORF3a and ORF6 interact with innate immune cellular proteins, indicating mechanisms of immunomodulation by the virus (27). Besides these mentioned genes, separate studies also showed that NSP2, NSP12, NSP16 and ORF8 involve immune response antagonism (28–32).

Overall, SARS-CoV-2 NSPs modulate host innate immune response at several levels by disrupting RNA splicing, translation, and protein trafficking (14, 16). The SARS-CoV-2 NSP1 deletion variant (Δ500-532), which became prevalent worldwide, is associated with a non-severe clinical illness and lower viral load, underlying the importance of NSP-mediated inhibition of IFN-I signaling (19). SARS-CoV-2 infection results in a shutdown of cellular protein translation via eIF2α which, at least in part, mediated by NSP1 (20), which also inhibits host gene expression throught disrupting mRNA nuclear export machinery and NSP1-binding to ribosomal channel (33, 34). A study also found that NSP1 prevents IFN-I induction, in part by blocking interferon regulatory factor 3 (IRF3) phosphorylation and NSP1-induced depletion of Tyk2 and, therefore, inhibiting the induction of IFN-I stimulated genes (ISGs) (21). NSP2 directly interacts with host GIGYF2 protein to enhance the interaction of GIGYF2 with 4EHP leading to inhibition of translation of Ifnb1 mRNA by GIGYF2/4EHP translational repressor complex (28). NSP3 contributes to the cleavage of ISG15 from IRF3 and attenuates IFN-I responses (38). NSP5 is known to target RIG-I and MAVS to evade the innate immune response, and NSP5 with mutation C145A failed to inhibit induction of IFN-I (39). NSP6 was reported to play roles in the suppression of IFN-I response in host cells through interacting TBK1 to suppress IRF3 phosphorylation (15) while NSP12 did not impair IRF3 phosphorylation but suppressed its nuclear translocation and ability to activate IFN promoter to attenuate IFN-I production (29). NSP13 was suggested to interact with the host kinase TBK1, potentially inhibiting IFN induction or directly target IRF3 to antagonize IFN-I response (15, 35, 36). NSP15 is a uridine specific endoribonuclease conserved across the *Coronaviridae* family (37) and mediates the evasion of host recognition of viral dsRNA (38). The coronavirus NSP14 and NSP16, which are N-7- and 2-Oʹ-methyltransferase respectively, add cap to the viral RNAs so that they can evade immune recognition (39). NSP14 was found to suppress IFN-β production, but the mechanism is still not fully clear (23, 25). NSP16 shields viral transcripts from detection by MDA5 and counteracts viral inhibition by IFIT1 (30). Replication of the virus with mutated NSP16 is highly sensitive to IFN-I, and the mutant triggers an enhanced MDA5-dependent immune response after viral infection (30).

SARS-CoV-2 accessory proteins also evolved strategies to antagonize IFN-I response (40). The ORF3a is the specific polymorphic protein of SARS-CoV and SARS-CoV-2 that activates the innate immune cells and induces cytokine storm (41). ORF3a blocks STAT1 phosphorylation to suppress IFN-I signaling (15). When compared to SARS-CoV, SARS-CoV-2 appeared to be more sensitive to IFN-I treatment, less efficient in suppressing cytokine induction via IRF3 nuclear translocation, and permissive of a higher level of induction MX1 and ISG56 (42). SARS-CoV-2 ORF6 expressed in the context of a fully replicating SARS-CoV backbone suppressed MX1 induction, but this suppression was less efficient than that by SARS-CoV ORF6 (42). Despite the sequence differences between SARS-CoV-2 and SARS-CoV ORF6, the antagonism of IFN-I signaling by SARS-CoV ORF6 was also observed with SARS-CoV-2 ORF6, as it inhibited IFN-β production at the level of or downstream of IRF3 activation (19, 20, 25, 35). ORF6s interact with transcription factors of the IFN-I pathway to antagonize the cytokine production (43–45). The ORF8 protein of SARS-CoV-2 is an Ig-like domain protein which has a limited similarity with its SARS-CoV counterpart and exhibits multiple immune-evasive functions (46). A recent study showed that deletion of ORF8 attenuated SARS-CoV-2 *in vitro* and in mice, with an increase in IFN-I induction and signaling (32).

The immune antagonism studies described above relied *in vitro* cell cultures transfected with plasmids over-expressing SARS-CoV-2 proteins to demonstrate their role in IFN-I antagonism (15, 18, 23, 25). However, this system most likely does not not recapitulate biological effects of the proteins in the context of authentic SARS-CoV-2 infection. For example, a study found that expression of ORF7a in transfected cells reduces surface expression of MHC-I (47). This conclusion has largely come from the reporter assays using co-transfection of cells with IFN-I signaling-related reporter construct(s) and plasmids over-expressing selected SARS-CoV-2 proteins. However, no effect was observed with an ORF7a-deleted replication-competent virus. Importantly, the previous studies characterized the effects of innate immune-antagonizing viral proteins individually and in different experimental systems, making assessment of their relative importance impossible.

Here we investigated inhibition of IFN-I signaling by viral proteins in the context of replication-competent SARS-CoV-2. We generated twelve infectious SARS-CoV-2 viruses with mutations in individual proteins and demonstrated that many of them have IFN-I-antagonizing activity and immunosuppressive effects in human cells and in the K18 hACE mouse model of infection. Our data indicates that SARS-CoV-2 antagonizes multiple immune mechanisms, particularly IFN-I signaling, activation of innate immune cells and T and B lymphocyte functions with the largest effects due to NSP1 and NSP15.

## RESULTS

### Generation of SARS-CoV-2 mutant viruses

To investigate the contribution of SARS-CoV-2 proteins or domains on viral pathogenesis and protective immune mechanisms, we selected 13 genes and specific mutations based on the available literature on several coronaviruses (15, 18, 21, 25, 28–31, 35, 36, 48–50) (Fig. 1A). To generate SARS-CoV-2 mutants, we first introduced mutations into the viral genes in an infectious cDNA clone pBeloBAC11-SARS-CoV-2 (51), which was driven by the human cytomegalovirus immediate early gene promoter, using *en passant* mutagenesis (52) (the primers for mutagenesis are shown in Suppl. Table 1). After the mutations were confirmed in the cDNA clones by Sanger DNA sequencing, Vero E6 cells were transfected with the cDNA clones containing the desired mutations for recombinant virus recovery. We generated a total of 16 mutated cDNA clones (Fig. 1A, B). To recover viruses from the mutated cDNA clones, 80% confluent monolayers of Vero E6 cells in 12-well plates were transfected with 1.0 μg per well of infectious SARS-CoV-2 BAC DNA WT or its mutated derivatives. At 48 h post transfection, transfected cells were split and seeded into T25 flasks with 50% confluent monolayers of Vero E6 cells. The cell cultures were observed daily until the appearance of cytopathic effect to collect viral supernatants. Viruses were recovered from the wild-type (WT) non-mutated control and the 12 cDNA clones containing mutations in NSP1, NSP2, NSP3, NSP6, NSP12, NSP13, NSP14, NSP15, NSP16, ORF3a, ORF6 or ORF8 (Fig. 1A). The recovered viruses were expanded by additional 1 or 2 passages, and the viral genomic RNA was isolated from the cell culture supernatants. The mutations in the viral genomes were then confirmed by sequencing (Suppl. Fig. 1). We could not recover viruses from the cDNA clones carrying two separate mutations in NSP3, or a mutation in NSP5, or a deletion in NSP6 (Fig. 1A). All the viruses were titrated in Vero E6 cells by plaque assay and their plaque sizes were compared. We found that all the mutant viruses, except the NSP2 and NSP3 mutants, have a smaller plaque phenotype when compared to WT virus (Fig. 1C).

**Fig. 1.**
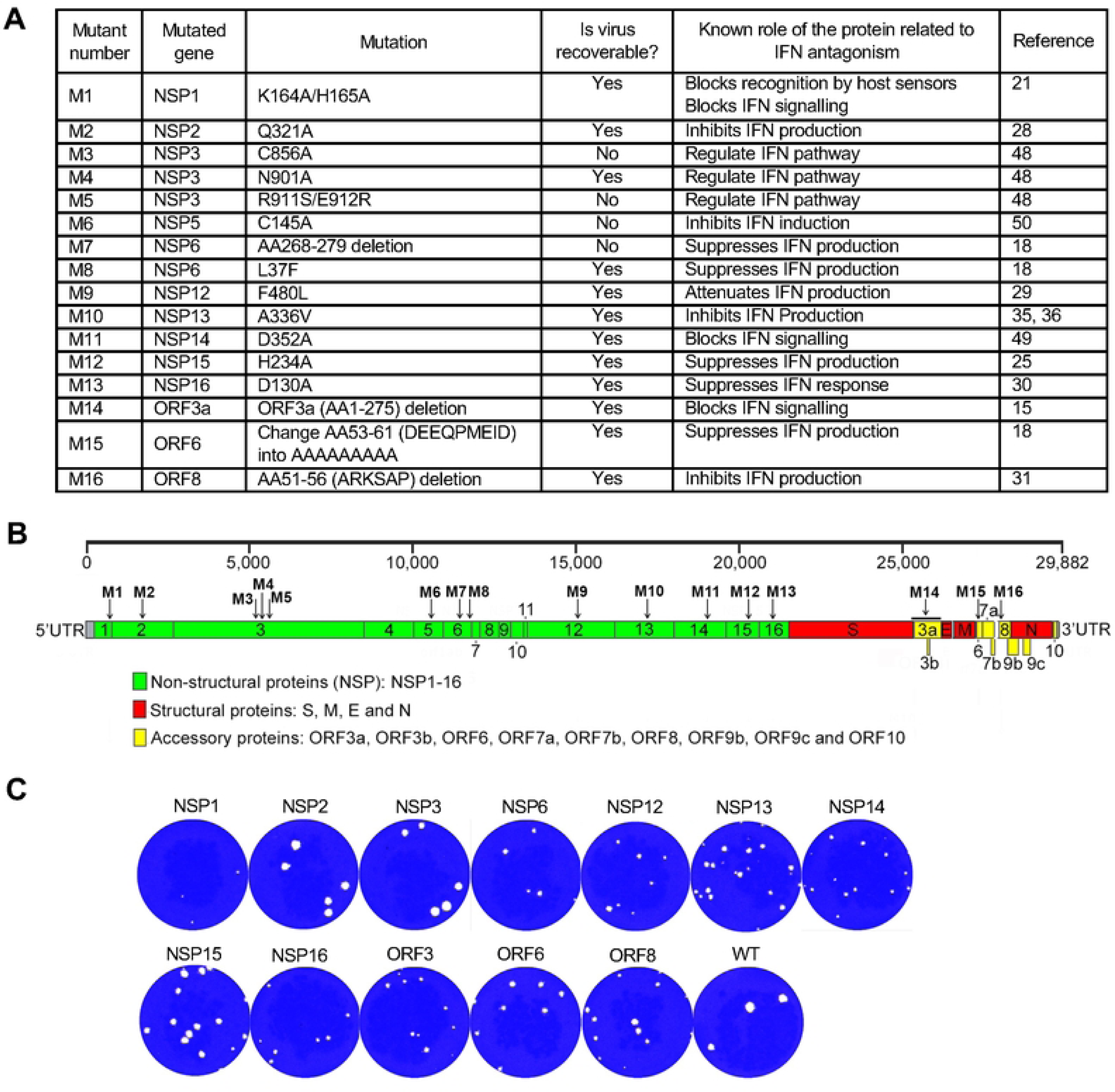
Generation of SARS-CoV-2 mutant strains. **A.** List of the mutations in the indicated viruses. **B.** Schematic representation of the SARS-CoV-2 genome. The positions of the mutations are indicated by arrows with mutation numbers. **C.** Comparison of plaques of WT SARS-CoV-2 and the mutants on day 2.

### Multiple SARS-CoV-2 NSP proteins contribute to inhibition of IFN-I induction

Using the panel of 12 mutated SARS-CoV-2, we investigated the impact of NSP and ORF proteins on IFN-I signaling. We transfected 293T-ACE2/TMPRSS2 cells, constitutively expressing SARS-CoV-2 receptors ACE-2 and transmembrane serine protease 2 (TMPRSS2) (ACE2/TMPRSS2) (53), with pISRE-luc firefly luciferase reporter plasmid and pTK-RL renilla luciferase reporter plasmid for control of transfection efficiency. Next day, cells were mock-infected or infected with SARS-CoV-2 WT or mutant viruses at 1.5 plaque-forming unites (PFU) per cell. After 24 h of infection, cells were either left untreated or treated with IFN-α (100 U/ml) and incubated for an additional 24 h, when ISRE-induced luciferase activity was measured (Fig. 2A). As expected, WT SARS-CoV-2 inhibited ISRE-luc activity when compared to mock-infected cells (Fig. 2B). Importantly, significantly higher ISRE-luc activities were observed in cells infected with NSP1, NSP15 and NSP16 mutant viruses as compared to WT SARS-CoV-2 (Fig. 2B). Even with IFN-α treatment, WT SARS-CoV-2 inhibited ISRE-luc activity (Fig. 2C). In contrast, most of the mutant viruses – NSP1, NSP2, NSP3, NSP6, NSP14, NSP15, ORF3a, ORF6 and ORF8 – shown higher ISRE-luc activity than WT SARS-CoV-2. These data suggest multiple and redundant mechanisms of IFN-I antagonism mediated by SARS-CoV-2, with the major effects mediated by NSP1, NSP15 and NSP16, and lesser effects mediated by the other viral proteins.

**Fig. 2.**
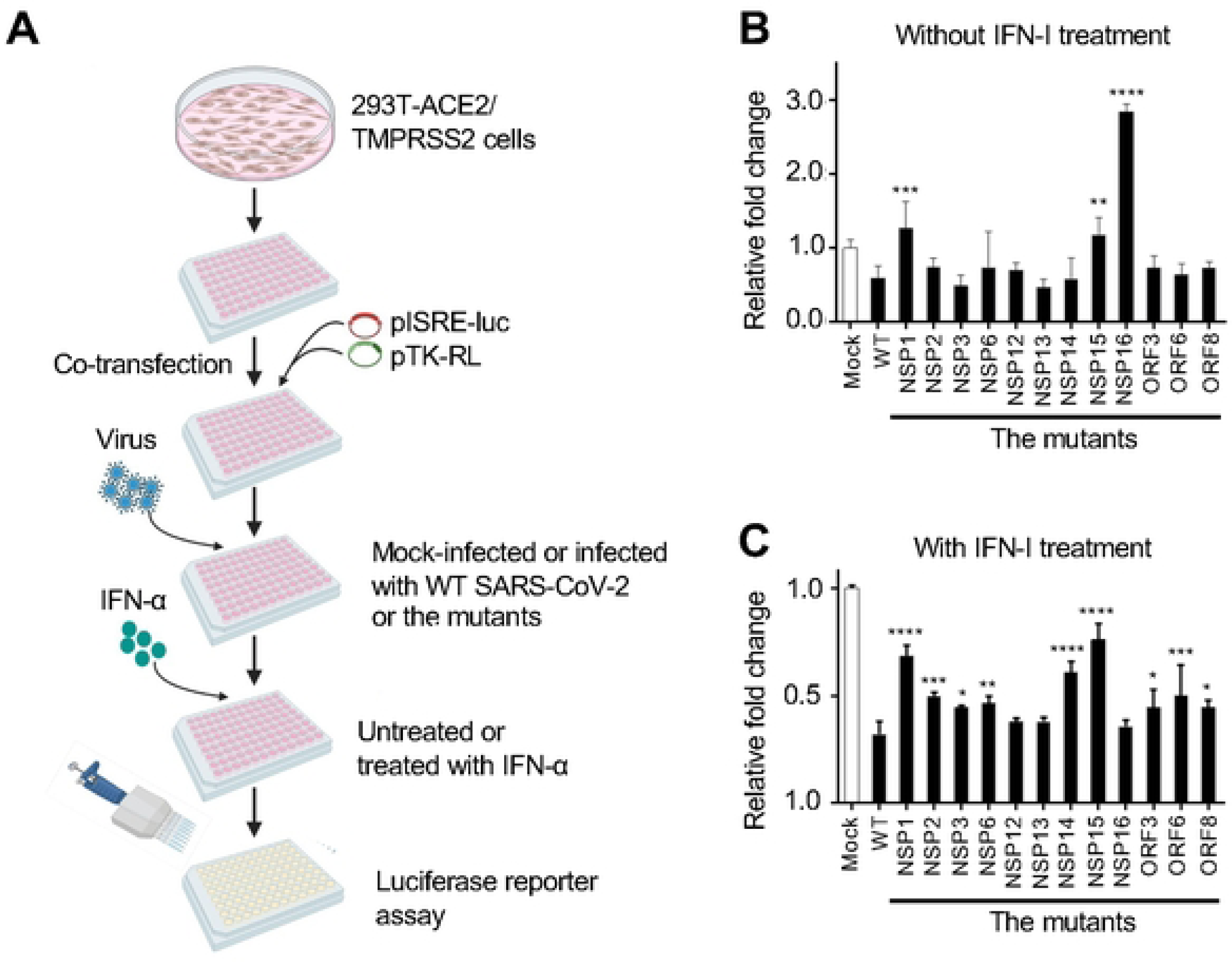
SARS-CoV-2 NSP1 and NSP15 are the most potent antagonists of IFN-l response. **A.** Flowchart of the experiment. 293T-ACE2/TMPRSS2 cells were transfected with plSRE-luc and pTK-RL At 24 h after the transfection the cells were mock-infected or infected with WT SARS-CoV-2 or the 12 mutants. At 24 h after the infection, cells were mock-treated or treated with IFN-α for additional 24 h, and the luciferase activity of thecell lysates was measured. **B, C.** Relative fold change in luciferase activity in 293T-ACE2/TMPRSS2 cells without (**B**) and with (**C**) IFN-α treatment. Mean values ± SD based on triplicate samples. The significance wasdetermined by one way ANOVA. * p < 0.05, ** p < 0.01, ***< 0.001, ****p<0.0001.

### SARS-CoV-2 NSP1 and NSP15 suppress maturation of human dendritic cells

SARS-CoV-2 modulates the function of T cells, natural killer cells, monocytes and specifically impairs the functional phenotypes of dendritic cells (DCs) (54–56). As noted above, SARS-CoV-2 antagonizes IFN-I production and signaling. IFN-I promotes the phenotypic and functional maturation of DCs (57–59). We therefore tested the effects of SARS-CoV-2 WT and mutants on the maturation of immature human monocyte-derived DCs. Cells were stimulated with LPS (positive control), SARS-CoV-2, WT or its mutants for 1, 2 or 3 days (Fig. 3A). The maturation status of CD11c+ DCs was assessed by flow cytometry measuring the expression of maturation markers CD80 and CD86, with HLA-DR/MHCII also assessed as a reference marker. As expected, LPS treatment significantly up-regulated expression of CD80 and CD86 at all three timepoints, as seen in mean fluorescent intensity (MFI) (Fig. 3B, C). However, SARS-CoV-2 showed no DC maturation based on the induction of the two markers, which was comparable to mock-infected DCs (Fig. 3C). Strikingly, the NSP1 and NSP15 mutant viruses induced the maturation of the two markers at a significantly higher level than WT SARS-CoV-2. In contrast, NSP13 reduced CD80 expression at day 3, which is a deviation from other mutants that did not alter the expression of the maturation markers. Although MHCII expression was noted in the majority of the cells, its MFI was highly variable depending on the stimulants (Fig. 3C). Together, these data indicate that SARS-CoV-2 suppresses DC maturation, and that the effect is mainly mediated by NSP1 and NSP15.

**Fig. 3.**
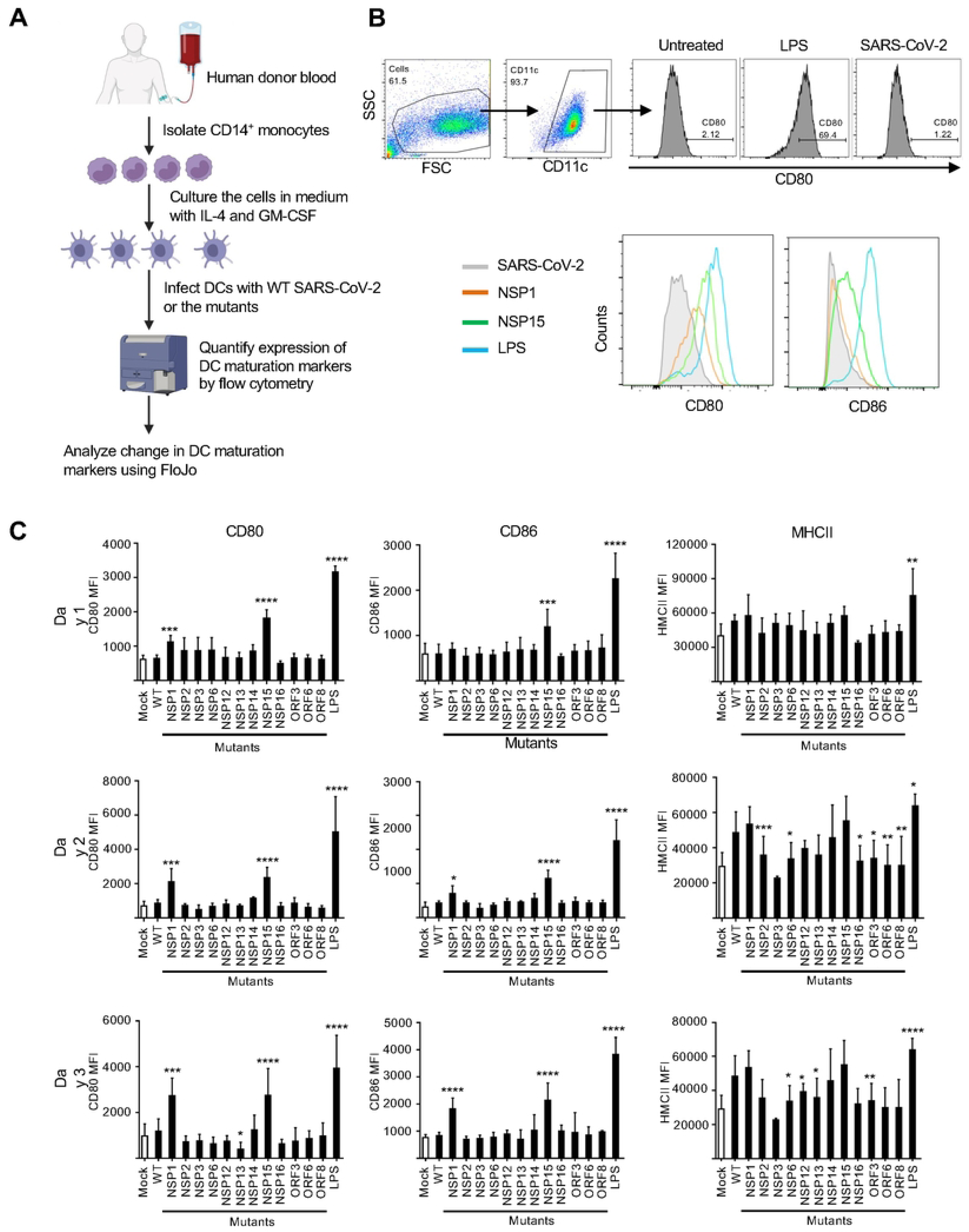
SARS-CoV-2 NSP1 and NSP15 suppress maturation of human dendritic cells. **A.** Flowchart of the experiment. CD14+ monocytes were isolated from fresh human PBMCs and further cultured in the presence of IL-4 and GM-CSF for differentiation into DCs. On day 6, the DCs were treated with LPS, or mock-infected or infected with WT SARS-CoV-2 or the 12 mutants at MOI of 3 PFU/cell, cultured for 1, 2 or 3 days and analyzed for expression of maturation markers by flow cytometry. **B.** Top: gating strategy with representative samples cultured for 2 days. Bottom: histograms showing expression of maturation markers CD80 and CD86 in representative samples cultured for 2 days. **C.** MFI of the DC maturation markers CD80, CD86 and also MHCII in CD11c+ DCs at day 1 (top panel), day 2 (middle panel) and day 3 (bottom panel). Data represent mean ± SD from three independent healthy donors in two technical replicates. The significance was determined by one way ANOVA: * p <0.05, ** p < 0.01, ***< 0.001, ****p<0.0001.

### SARS-CoV-2 NSP1 and NSP15 suppress genes that control innate immune response in Calu-3 cells

To explore the effects of SARS-CoV-2 proteins on expression of host genes, we first mock-infected or infected Calu-3 cells with SARS-CoV-2 WT or the 12 mutants. At 24 h post-infection the cells were used for isolation of RNA which was deep sequenced (Fig. 4A). Principal component analysis (PCA) of transcriptome from 42 samples (3 repeats for each virus and mock infections) demonstrated that gene expression patterns in the cells infected with SARS-CoV-2 WT and the 12 mutants were different from those in mock-infected cells and from each other (Fig. 4B). The differential expression analysis demonstrated that the mutants up-or down-regulated cellular gene expression to a different extent. NSP3, NSP13, NSP15 mutants upregulated more genes, while the NSP6, NSP14 and ORF3a mutants down-regulated more genes when compared to the WT virus. However, the NSP1 and ORF6 mutants both were able to up and down-regulated quite different sets of host genes (Fig. 4C).

**Fig. 4.**
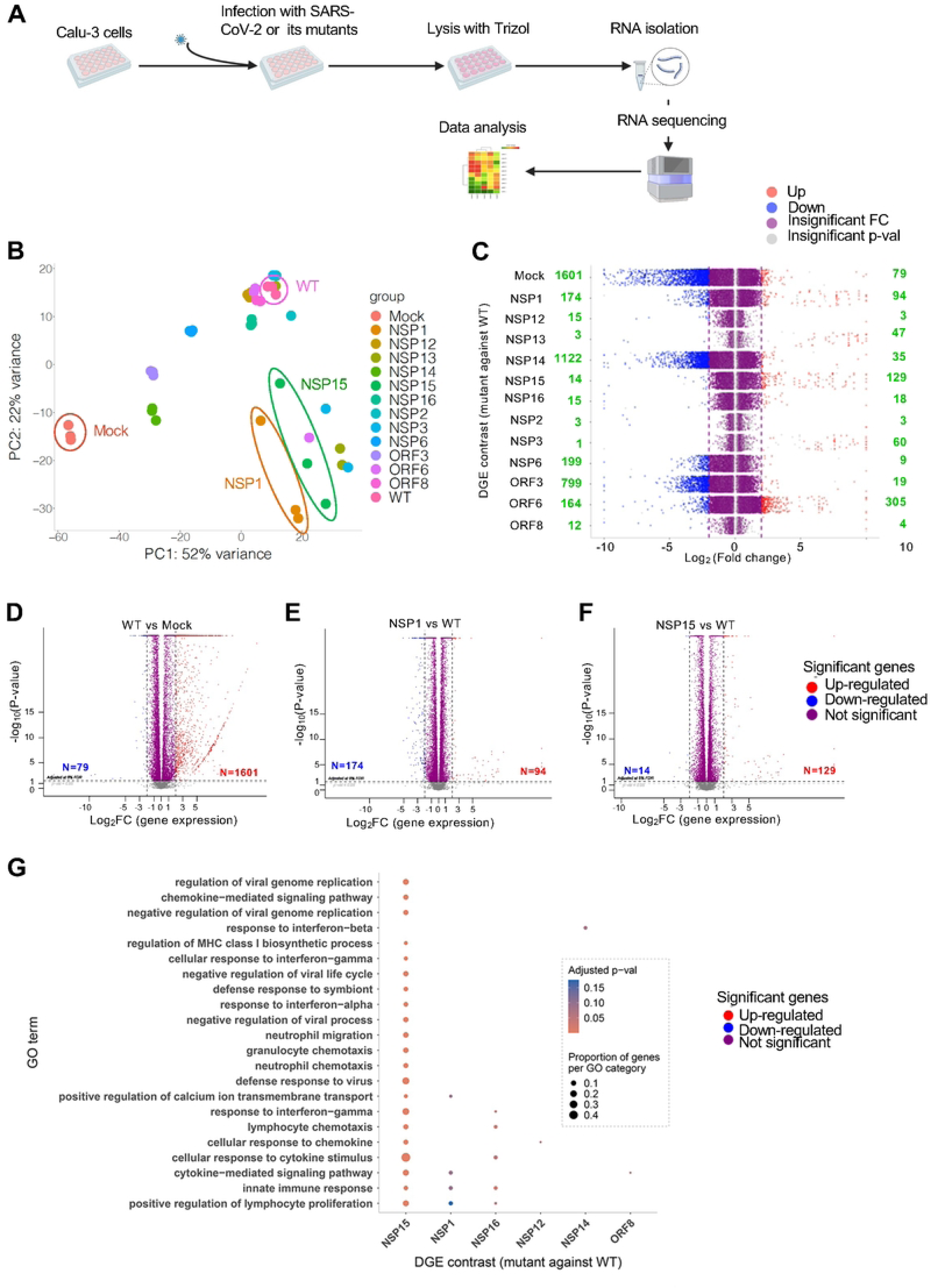
SARS-CoV-2 NSP1 and NSP15 suppress expression of genes involved in mechanisms of cell defense. **A.** Flowchart of the experiment. Calu-3 cells were mock-infected or infected in triplicates with *WT* SARS-CoV-2 or the 12 mutants, and 24 h later the cells were harvested in Trizol for RNA isolation, followed by RNA sequencing and data analysis. **B.** Principal component analysis (PCA) to identify the distance or clusters of gene activation by *WT* SARS-CoV-2 or the 12 mutants. **C.** Jitter plot analysis for change in differential gene expression (OGE) in Calu-3 cells mock-infected or infected with the 12 SARS-CoV-2 mutants: comparison *WT* SARS-CoV-2-infected cells. The numbers of up-and down-regulated genes are indicated in green. **D.** Volcano plot for OGE in *WT* versus mock. 1,601 genes were up-regulated (L092FC >2) and 79 genes were down­regulated (Log,FC >2) in *WT* SARS-CoV-2-infectedCalu-3 cells compared with mock-infected cells. **E.** Volcano plot for OGE in NSP1 versus WT. 94 genes were up-regulated (Log_2_FC >2) and 174 genes were down­ regulated (Log_2_FC >2) in NSP1 mutant-infectedCalu-3 cells compared with mock-infected cells. **F.** Volcano plot for OGE in NSP15 versus WT. 129 genes were up-regulated (Log_2_FC >2) and 14 genes were down­regulated (Log_2_FC >2) in NSP15 mutant-infected Calu-3 cells compared with mock-infected cells. **G.** Gene Ontology (GO) tenm analysis showing pathway enrichment for NSP1 and NSP15 over WT. NSP1, NSP15 inhibit several pathways including innate immune response and cy1okine-mediate signaling pathways.

Infection of Calu-3 cells with WT SARS-CoV-2 up-regulated more than 1,500 genes compared to mock-infected cells (Fig. 4D). The upregulated genes included those involved in innate immune response and are known responders of IFN-I stimulation (60): *IFNB1, CXCL10, OAS2, TNFAIP3 RSAD2, IFNL2, USP18, MX1, CXCL11, NFKBIA, IFIT2, ATF3, IFIT1, IFNL3,* and *IFI44L*. Among these, OAS2 is an important antiviral protein involved in innate immune response against viruses including SARS-CoV-2 (61). IFIT2 (ISG54) is an anti-viral protein that inhibits expression of viral mRNA and acts as a mediator of cell apoptosis (62). Infection with the NSP1, NSP12, NSP13, NSP14, NSP15, NSP16, NSP2, NSP3, NSP6, ORF3a, ORF6 or ORF8 mutant resulted in 94, 3, 47, 35, 129, 18, 3, 60, 9, 19, 305, and 4 cellular genes, respectively, upregulated (log_2_FC ≥2 and p-value <0.05) and 174, 15, 3, 1122, 14, 15, 3, 1, 199, 799, 164, and 12 genes, respectively, down-regulated (log_2_FC <2 and p-value <0.05) compared to that of WT SARS-CoV-2-infected cells (Fig. 4C-F).

Gene ontology (GO) analysis of the upregulated genes in the Calu-3 cells infected with these mutants revealed a significant enrichment in immune and inflammatory processes with different degrees (Fig. 4G, Suppl. Fig. 2). Although NSP1, NSP3, NSP13, NSP15 and ORF6 mutants upregulated different sets of genes in Calu-3 cells, NSP1 and especially NSP15 mutants have uniquely activated genes that are related to the inflammatory processes and cytokine responses (Fig. 4G). For example, for the NSP15 mutant, the upregulated pathways included cell response to cytokine signaling, defense response to virus, response to IFN-γ, cytokine-mediated signaling pathway, positive regulation of lymphocyte proliferation and a few others (Fig. 4G). Many of the genes upregulated by the NSP15 mutant are ISGs and are involved in innate defense against viruses. For example, the IFITMs are IFN-I-induced anti-viral proteins which inhibit SARS-CoV-2 replication (63), but sometimes promote it (64). *IFNL1* to *3* are primarily expressed in epithelial cells with major anti-viral properties (42, 65). The NSP1 and NSP15 mutant viruses also activated many chemokine and pro-inflammatory cytokine genes that could play a protective role during infection. While pro-inflammatory genes (CCL22, CCL5, CX3CL1, CXCL9-11, IL-1B, MMP1, TNFSF13 and 15) facilitate inflammation, the immune regulatory genes (CIITA, SSA1 and CFB) could control excessive inflammation (66–68). Importantly, IL-1β is important for adaptive immune cell differentiation and protective memory response (69, 70).

### SARS-COV-2 NSP1 and NSP15 contribute the viral pathogenicity in mice

We have shown that NSP1 and NSP15 suppress IFN-I signaling (Fig. 2) and DC maturation (Fig. 3). To investigate the impact of NSP1 and NSP15 on survival and gene expression in context of *in vivo* infection, we used the K18 hACE2 transgenic mouse model of COVID-19 (71). We performed three independent mouse infection experiments. In Experiment 1, we compared the clinical response of mouse (body weight change and lethality) following infection with WT SARS-CoV-2 or the NSP1 and NSP15 mutants (Fig. 5A-C). In Experiment 2, infected mice were euthanized on days 2 and 4 post-infection for assessment of the viral load in nasal turbinates and lungs (Fig. 5D,E), lung pathology (Fig. 5F-I) and transcriptome by bulk RNA-seq (Fig. 6). In Experiment 3, infected mice were euthanized on day 3 to collect lung cells for scRNA seq analysis (Fig. 7). In all the experiments, mice were infected intranasally with 10^5^ PFU of WT SARS-CoV-2 or NSP1 or NSP15 mutant. All the mice infected with WT SARS-CoV-2 succumbed to infection on days 6 – 8 (100% lethality), while the NSP1 and NSP15 mutants demonstrated 75% and 25% lethality, respectively (Fig. 5B). Consistent with the lethality pattern, mice infected with WT SARS-CoV-2 lost 25% of their initial body weight, while mice infected with SARS-CoV-2 NSP1 and NSP15 mutants lost 15% and 17%, respectively, of their initial body weight (Fig. 5C). Consistently with previous studies (72), WT SARS-CoV-2-infected mice demonstrated a very high viral load, both in the nasal turbinates and in the lungs (Fig. 5D). NSP1 mutant virus-infected mice had significantly lower viral load in the nasal turbinates on day 2 and in the lungs on days 2 and 4 post-infection compared to WT virus-infected mice. Consistently with the higher percentage of survival, mutant virus-infected mice had lower viral load in the nasal turbinates on day 2 and in the lungs on day 4, compared to WT virus-infected mice (Fig. 5E).

**Fig. 5.**
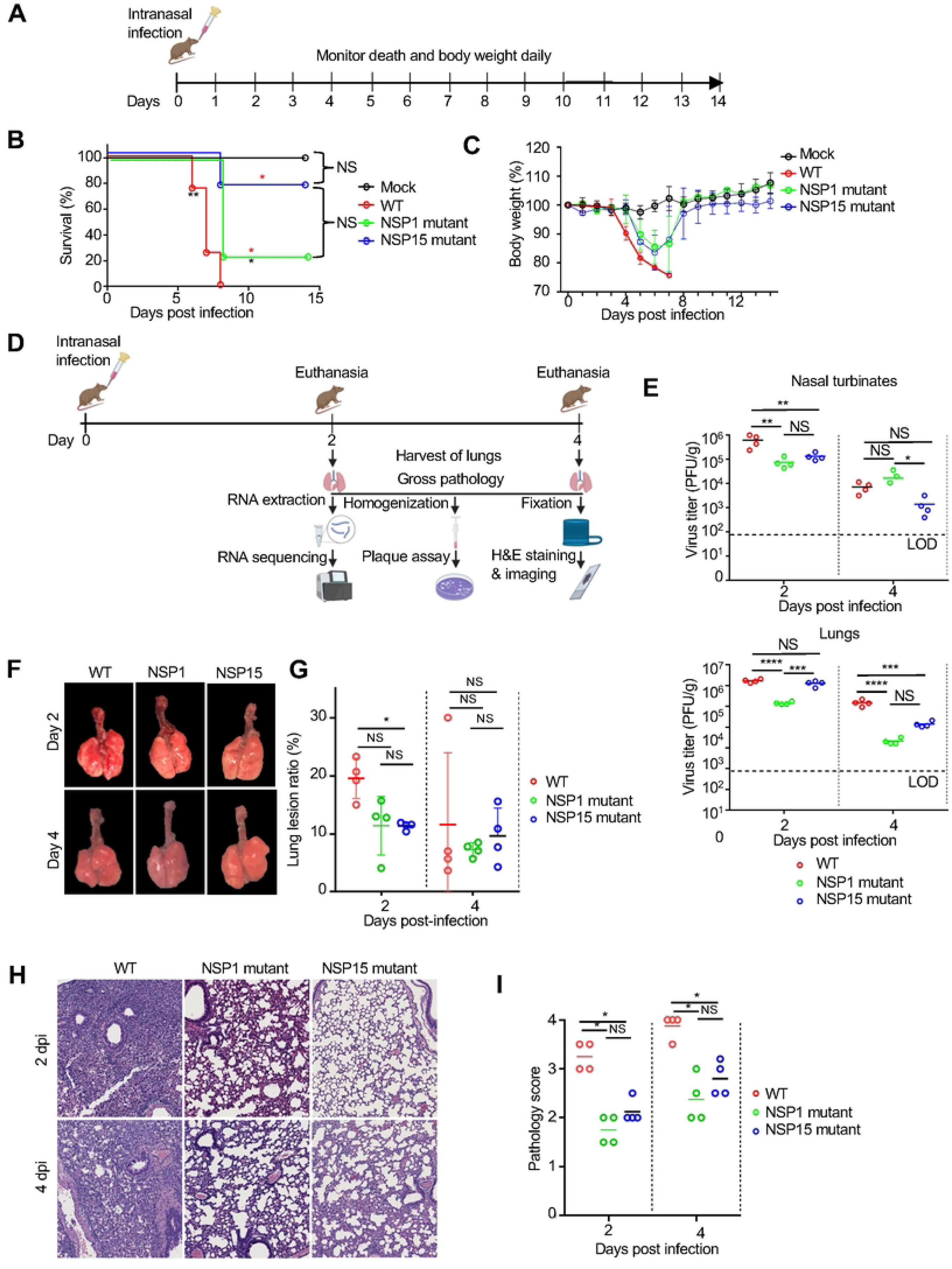
SARS-COV-2 NSP1 and NSP15 contribute the lethality and lung pathology. **A.** Experimental scheme of the mouse experiment 1 (panels A-C). Four groups of K18-hACE2 mice (n=4 per group) were mock-infected or infected intranasally with WT SARS-CoV-2 or NSP1 or NSP15 mutant at 1×10^5^ PFU. **B.** Survival. The red start(s) and black start are analysis results from the indicated virus compared to WT and mock respectively. **C.** Body weight. **D.** Experimental scheme of the mouse experiment 2 (panels D -I). Eight groups of K18-hACE2 mice (two groups for each virus or mock infection, n=4 per group) were mock-infected or infected intranasally with WT SARS-CoV-2 or NSP1 or NSP15 mutants at 1×10^5^ PFU, then euthanized at 2 or 4 days after the infection to harvest the lungs and nasal turbinates. **E.** Viral load in the nasal turbinates (top) and lungs (bottom). **F.** Lung images and pathology scores. **G.** Pathology scores. **H.** Lung histology: H&E staining. **I.** Histopathology score. Data shown in panels E, G, I represent mean ± SD from 4 animals. The significance was detennined by log-rank (Mantel­ Cox) test in panel B, one way ANOVA test in panel E, and Mann Whitney test in panels G, I. * p <0.05, **p < 0.01, *** < 0.001, ****p<0.0001.

**Fig. 6.**
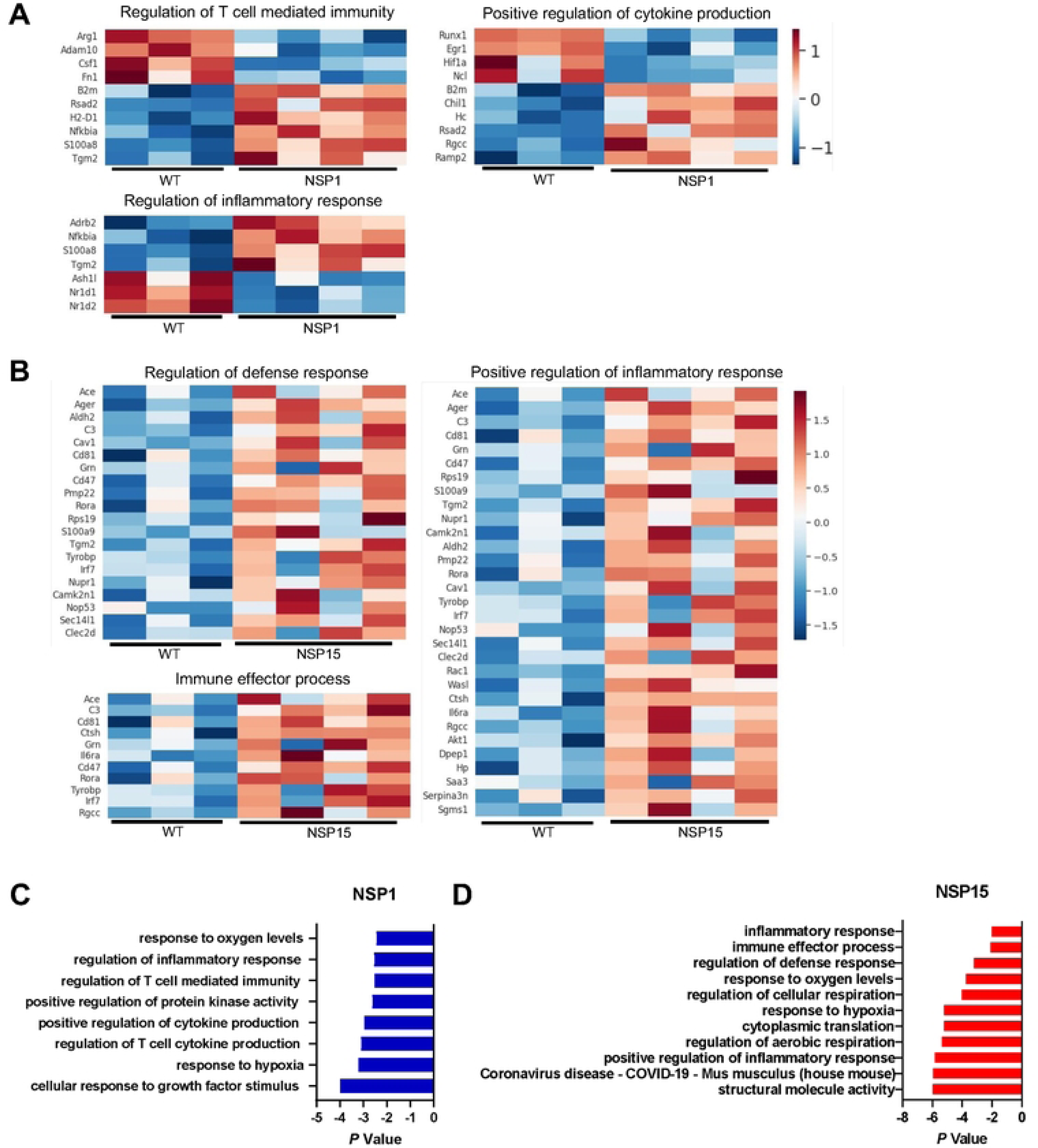
Bulk RNA-seq for transcriptome analysis of lungs in hACE2 transgenic mice infected by SARS-CoV-2 NSP1 and NSP15 mutants on day 2. Data from the mouse experiment 2 (Fig. 5D). **A.** Changes in selected pathways for the NSP1 mutant vs WT: regulation of T cell mediated immunity, positive regulation of cytokine production and regulation of inflammatory response. **B.** Changes in selected pathways for the NSP15 mutant vs WT: regulation of defense response, immune effector processes and positive regulation of the inflammatory response. **C.** Pathways upregulated for the NSP1 mutant vs. WT. **D.** Pathways upregulated for the NSP15 mutant vs WT.

**Fig. 7.**
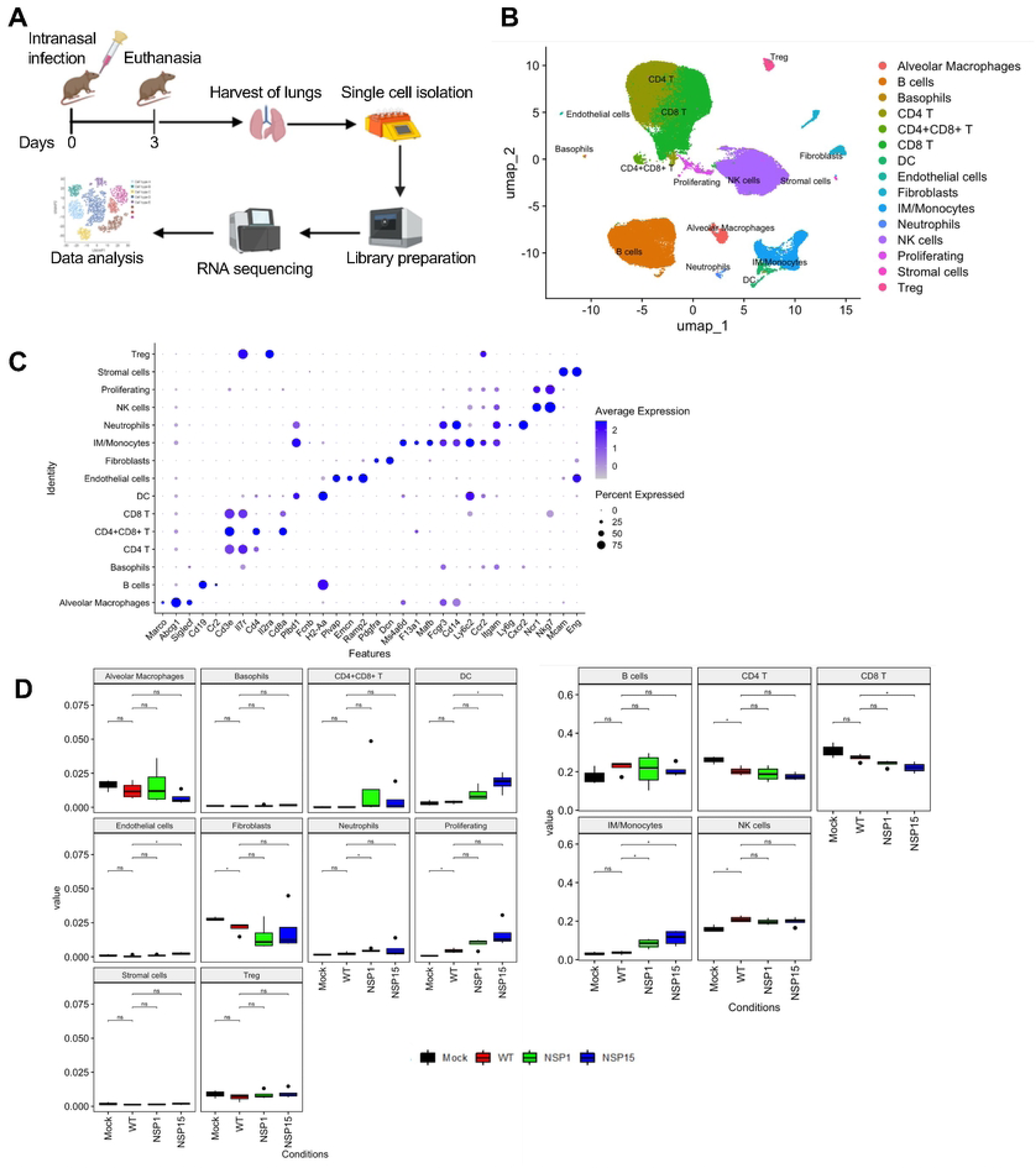
Single ell RNA-seq analysis of pulmonary tissues identifies cellpopulation differences between animals infected with WT, NSP1 or NSP15 mutant virus. **A.** Experimental scheme of the mouse experiment 3. **B.** Integrated UMAP (uniform manifold approximation and projection) of all Mock (n = 3), WT (n = 4), NSP1 (n = 4), and NSP15(n = 4) samples. **C.** Dot plot showing canonical marker expression in observed lung cell types. **D.** Box plots comparing cell-type proportions observed in Mock, WT, NSP1, and NSP15 groups. The limits of the box reflect the interquartile range (IQR: 03-01) with median shown as horizontal bars. Whiskers extend to 1.5 times the IQR of the box. For each cell type, pairwise I-test comparisons of the wild type (WT) proportion with every other group are shown(* Holm p-adj < 0.1, ns - non-significant).

Pathological examination of the lung tissues on day 2 showed that mice infected with WT SARS-CoV-2 show areas of lung congestion and consolidation. These changes were minimal in mice infected with the NSP1 and NSP15 mutants (Fig. 5F, G). Furthermore, mice infected with NSP1 or NSP15 mutants had reduced gross pathology compared to WT SARS-CoV-2-infected mice. However, on day 4, the pathological changes in mice infected with NSP1 or NSP15 mutants were comparable to that in WT-infected mice (Fig. 5F, G). On histological analysis, a typical interstitial pneumonia was noticed in lungs from WT SARS-CoV-2 infected mice on days 2 and 4 post-infection. The interstitial pneumonia was characterized by a severe inflammation with mononuclear cells and neutrophils in the alveolar space, interstitial septa and airways septal thickening, perivascular cuffing and vascular endothelial cell damage. In contrast, in mice infected with the NSP1 or NSP15 mutant, inflammatory changes are tempered with mild to moderate inflammation and minimal vascular and airway changes (Fig. 5H). Histopathology scores of lung sections (Fig. 5I and Suppl. Table 2) indicated that the NSP1 and NSP15 mutants caused less tissue damage than WT SARS-CoV-2 in the lungs.

Altogether, these data indicate that SARS-CoV-2 NSP1 and NSP15 suppress the immune mechanisms which contribute to the protection against viral disease and death.

### SARS-COV-2 NSP1 and NSP15 suppress innate immune responses in the lungs

As NSP1 and NSP15 contribute to viral pathogenicity in mice, we next investigated molecular pathways affected by the viral proteins in the lungs by comparison of transcriptome profiles in the lungs on day 4 by bulk RNA-seq analysis (mouse experiment 3). The DESeq2 analysis demonstrated 2,448 upregulated genes (log2 FC ≥2 and p-value <0.01) in lungs of mice infected with WT SARS-CoV-2 compared to mock-infected mice. On comparison with mice infected with WT SARS-CoV-2, there were 63 and 22 up-regulated genes [log2 fold change (FC) ≥2 and p-value <0.01] and 45 and 25 down-regulated genes (log2 FC ≥-2, p-value <0.01) in lungs of mice infected with SARS-CoV-2 NSP1 and NSP15 mutants, respectively (Suppl. Fig. 3). Many of the genes upregulated by the two mutants are associated with effector immune functions, including Ace*, C3, Ctsh, Rora, Irf7, Rgcc, Saa3, Grn,* and *Dpep7.* Some of the downregulated genes including *Arg1*, *Adam10*, *Robo2*, and *Smad2* are related to negative regulation of immune responses, suggesting that mutant viruses have activated the genes involved in effector immune functions. The heatmaps of DEG showed a quite different pattern of gene expression between the NSP1 and NSP15 mutants on one hand and WT virus on the other hand (Fig. 6A, B and Suppl. Fig. 4). The NSP1 and NSP15 mutants up-regulated, as compared to WT, expression of 534 and 242 genes, respectively. In NSP1 mutant-infected mice upregulated genes included those involved in regulation of T cell immunity (B2m, h2d1, nfkbia, s100a8 and tgm2) and regulation of cytokines/inflammatory response (*b2m*, *nfkbia*, and *ramp2*) (Fig. 6A) (73–77). In NSP15 mutant-infected mice, up-regulated genes included those involved in immune effector and inflammatory processes, particularly of innate immunity – *C3, CD81, Ctsh, Grn, Il6ra, Rora, Tyrobp, Irf7, Rgcc, Kamk2n1, Cav1, Dpep1*, *Hp*, and *Saa3* (Fig. 6B). Products of *C3, Ctsh, Grn, Il6ra*, and *Rora* genes are critical for functions of innate immune cells (78–82). *Cd81* is important for B cell differentiation and proliferation (83), while *Cd47* is an integrin-associated protein important for transmigration of leukocytes (84). *Tyrobp, Kamk2n1, Cav1,* and *Dpep1* regulate several protein kinases involved in immune cell function, particularly SYK, ERK1/2, MAPK and PI3K/Akt/mTOR. IRF7 plays a vital role in the transcriptional activation of virus-inducible cellular genes and particularly IFN-I genes (85). The *Saa3* and *Hp* genes encode acute phase proteins with pro-inflammatory and immunostimulatory effects (86, 87). Peroducts of many of these genes are involved in immune effector functions. Importantly, the NSP1 and NSP15 mutants also down-regulate, as compared to WT, expression of certain genes, particularly *Runx1, Arg1, Junb, Cox5a, Adam10, Hif1a,* and *Serbp1*, that are transcriptional repressors and are involved in excessive inflammatory responses through hematopoiesis.

Pathway enrichment analysis revealed activation of genes in several pathways that are related to immune responses and other cellular functions for both the NSP1 and NSP15 mutants (Fig 6C, D). However, the changes in the pathway profiles induced by the two mutants were different. Together, bulk RNA-seq analysis revealed that NSP1 and NSP15 are involved in suppression and perturbation of immune effector functions that confer protective immunity against SARS-CoV-2.

### SARS-CoV-2 NSP1 and NSP15 suppress activation of immune functions in the mouse lung

To elucidate the effects of SARS-CoV-2 NSP1 and NSP15 on lung immune cells, we performed single cell RNA-seq analysis of cells isolated from the lungs of K18 hACE2 transgenic mice on day 3 after infection with either WT SARS-CoV-2, the NSP1 or the NSP15 mutant or after mock-infection (Fig 7A). We identified 15 major cell UMAP clusters: B cells, basophils, CD4 T cells, CD4+CD8+ T cells, CD8 T cells, DCs, endothelial cells, fibroblasts, alveolar macrophages, monocytes, neutrophils, NK cells, proliferating, stromal cells, and Tregs (Fig 7B-C). The alveolar macrophage population exhibited markers typical for these cells, including *Marco, Abcg1 and Siglecf* (Fig 7C, Suppl. Fig. 5). The monocyte cluster was an intermixed population of cells that included both monocytes (marked by *Fcgr3, Cd14, Ly6c2, Ccr2,* see Fig 7C, Suppl. Fig. 5) and interstitial macrophages (marked by *Ms4a6d, F13a1, Mafb,* see Fig 7C, Suppl. Fig. 5). This co-clustering is consistent with the close relationship of interstitial macrophages and monocytes (88, 89). We refer to this cluster as interstitial macrophage (IM)/monocytes.

In comparison with mock infection, WT infection resulted in a significant increase in the proportion of NK and proliferating cells and a significant reduction in the proportion of CD4+ T cells and fibroblasts. We also observed a reduction in the proportion of alveolar macrophages and CD8+ T cells that did not reach significance. These changes support immunosuppression by the WT virus. Infection with the NSP1 and NSP15 mutants resulted in an increase in the proportion of IM/monocytes as compared to WT-infected mice. Moreover, the relative expression levels of some monocyte and IM markers were differentially affected by WT and mutant infections (Suppl. Fig. 6). For example, the microenvironment modulatory IM marker C5ar1 is dramatically increased only after infection with the NSP15 mutant (Suppl. Fig. 6). These results indicate that NSP1 and NSP15 have specific effects on the state of IM/monocytes. The proportion of double positive CD4+/CD8+ T cells were increased with the mutant infections as compared to mock-infected and WT virus-infected mice. In addition, infection with the NSP15 mutant also resulted in an increase in the proportion of DCs and reduction of CD8+ T cells; similar trends were observed in infection with the NSP1 mutant without reaching statistical significance (Fig. 7D). Analysis of the viral transcripts demonstrated that SARS-CoV-2 viral reads (N or ORF1AB) were detectable in monocyte and macrophage populations, and also in B cells, but the levels of expression were inconsistent across the groups (Suppl. Fig. 7). Overall, these data suggest that NSP1 and especially NSP15 contribute to the immunosuppression and dysregulation of the immune response caused by SARS-CoV-2.

Differential gene expression analysis of WT virus infected mice compared to mock showed substantial regulatory remodeling of multiple cell types (Suppl. Fig 8A). In particular, there were 242 upregulated and 109 downregulated genes in IM/monocytes. Additionally, in all cell types analyzed other than proliferating cells, WT virus infection resulted in a regulatory program that included many more upregulated genes compared to downregulated genes (Suppl. Fig. 8A, Suppl. Table 3) . Pathway analysis revealed enrichment of several pathways linked to immune effector functions, primarily the innate immunity and the cellular metabolic pathways in response to WT virus infection, in all analyzed cell populations, with a somewhat lesser number of activated pathways detected in DCs and neutrophils (Suppl. Fig. 9). Alterations in the regulatory program between the mutants and WT virus were also observed (Suppl. Fig 8B,C). In comparison to WT virus, the NSP1 mutant showed upregulation of IFN-I response in IM/monocytes and T cell populations (Fig. 8A). In addition, the mutant showed downregulation of nucleoside-binding oligomerization domain (NOD) genes for some lymphoid cell populations. The NSP15 mutant, on the other hand, activated multiple pathways in IM/monocytes, but down-regulated several pathways in NK cells (Fig. 8B). Together, these data suggest that both NSP1 and NSP15 mutants suppress multiple pathways associated with antiviral defense, but the effects of these two proteins are different.

**Fig. 8.**
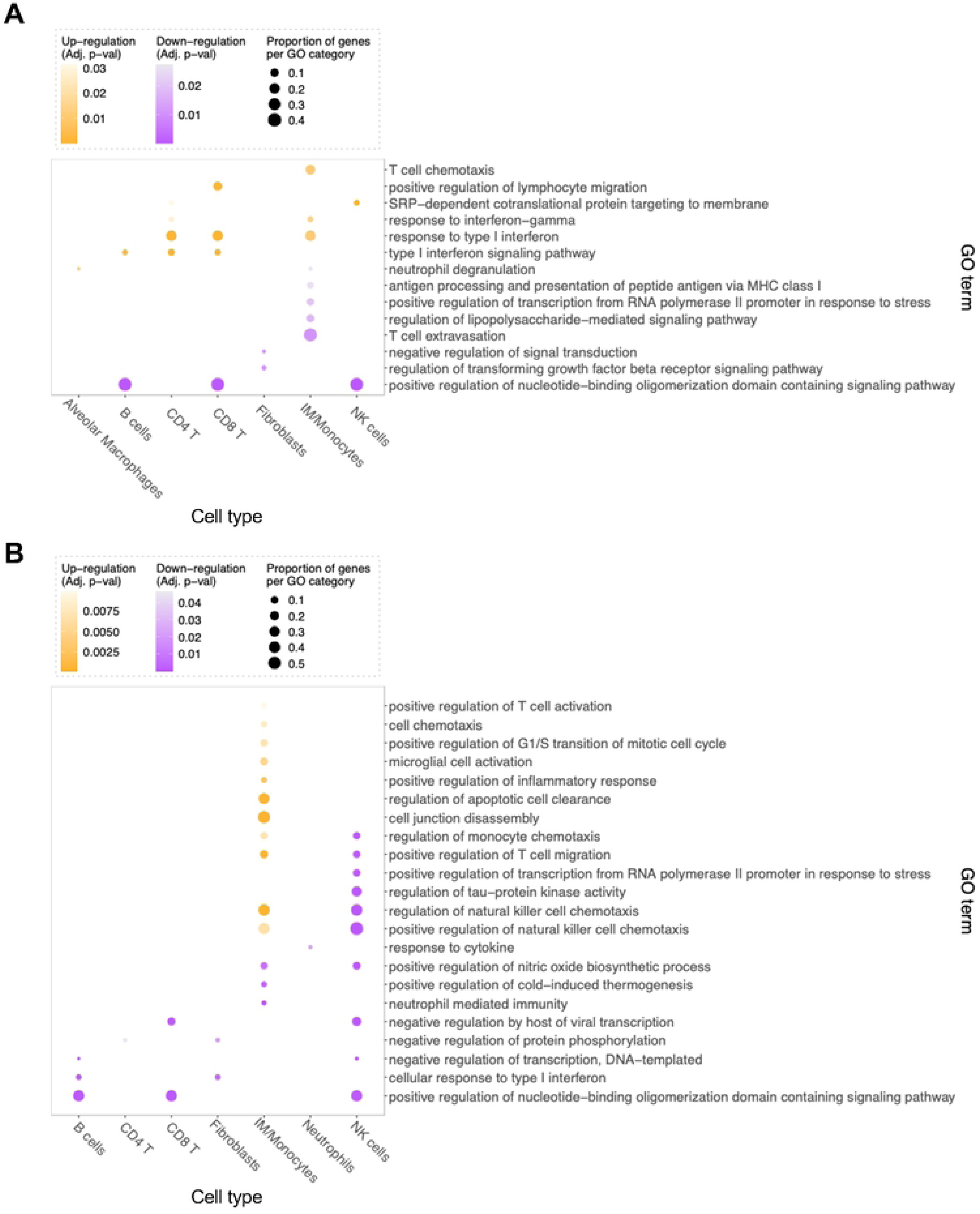
Change in activation of pathways associated with NSP1 and NSP15 mutants. Dot plots showing GO terms from pathway enrichment analyses by EnrichR (110) among differentially expressed genes (DEG) for the NSP1 vs WT (**A**), NSP15 vs WT (**B**). Dot size represents the fraction of DEG within the GO term. Dot color represents thedirection of the regulation of the term in the corresponding cell type (Up-regulation: yellow; Down-regulation: purple) and the color scale indicates the adjusted p-value (shown are only terms with FDR adjusted enrichment p-value < 0.1). Terms with substantial gene overlap are filtered out, with terms remaining only if there is a difference of at least two regulated genes from every other term. The direction of regulation for each enriched term is determined by the proportion of upregulated DEG versus thedownregulated DEG across all cell types.

## DISCUSSION

Induction of IFN-I and host innate immune responses provide a critical first line of defense against a wide variety of virus infections (90). Upon viral infection, viral RNA or DNA in the host cells triggers a signaling cascade to produce IFN-I (91). Then, IFN-I stimulates the expression of ISGs and other anti-viral genes through binding to surface receptors on the membranes of host cells to inhibit virus replication or/and prevent bystander uninfected cells from infection (92, 93). Like the other highly pathogenic coronaviruses SARS-CoV and Middle East respiratory syndrome coronavirus (MERS-CoV), SARS-CoV-2 evolved multiple strategies to evade IFN-I response (94) and, as a consequence, replicates to high titers in host tissues (95). Here, we focused on side-by-side comparison of multiple SARS-CoV-2 proteins which antagonizes the innate immune response. We selected multiple genes of SARS-CoV-2 based on their significance for viral pathogenicity and inhibition of the host immune responses to determine: (1) their contribution in immunosuppression in the context of viral infection, rather than a plasmid-based system, and (2) to compare their relative contribution to immunosuppression relative to each other side-by-side in multiple experimental systems. The studies showed that all the 13 genes play a role in IFN-I antagonism (Fig. 1A). Based on the information, we generated 16 full-length cDNA BAC clones with mutations in the 13 different genes (Fig. 1A). Among them, 12 clones resulted in a successful recovery of infectious viruses, while no virus could be recovered from the other four clones (Fig. 1A).

These irrecoverable constructs included two clones with mutations in NSP3 (R911S/E912R located close to the ISG binding site, and C856A in the catalytic site), one clone with mutation at C145A in NSP5 and one clone with mutation at NSP6 by replacing AA53-61 (DEEQPMEID) with AAAAAAAAA. NSP3 has the PLpro activity for NSP1-4 production, and also cleaves ubiquitin-like ISG15 protein from IRF-3 (48). NSP3 with the mutations R911S/E912R has reduced ISG-15 cleavage activity. Whether the two constructs with mutations in NSP3 were not recoverable due to the loss of PLpro and/or deubiquitinating activity, needs to be further investigated. Interestingly, we were able to recover virus from the clone with mutation N901A in the ISG-15 binding residue in NSP3, and the mutant virus formed larger plaques compared to WT virus (Fig. 1C). The mutant virus with deleted macrodomain (Mac1) in NSP3 induced increased levels of IFN-I and ISG expression in cell culture and mice (96). The Mac1 is located far away from N901 in NSP3. The mutant with the mutation N901A in NSP3 could induce a higher ISRE-luciferase activity after IFN-I treatment (Fig. 2C). This indicates that NSP3 has redundant strategies to inhibit IFN-I response to support viral replication. As NSP5 with the mutation C145A fails to inhibit induction of IFN-I (50), we also introduced this mutation in the full-length clone. However, we were not able to recover virus from the construct, presumably due to the loss of the 3CLpro activity (97). NSP3, NSP4 and NSP6 form replication-organelle-like structures which are required for SARS-CoV-2 replication (98). NSP6 is a membrane protein with predicted 7 transmembrane domains (98). So far, very little work has been done to map the functional domains of NSP6. The fragment AA269-279, which we deleted, is located downstream of the seventh predicted transmembrane domain and very close to the C-terminus of the protein. We couldn’t recover the virus from the construct with the AA269-279 deletion in NSP6. We speculate that the deletion may disrupt the formation of replication-organelle-like structures.

We used the ISRE-driven luciferase assay in 293T-ACE2/TMPRSS2 cells to first test the 12 mutated viruses without and without IFN-α treatment. Remarkably, seven of them demonstrated induction of IFN-I response in at least one of the two systems (Fig. 2). In IFN-α-treated cells, the NSP1 and NSP15 mutant viruses demonstrated the greatest response. NSP1 interacts the human 40S to interfere with mRNA binding and translation. SARS-CoV-2 NSP1 with mutations K164A/H165A lost its ribosome binding and inhibition capability (34). NSP15 is a uridine specific endoribonuclease conserved across the *Coronaviridae* family (37) and mediates the evasion of host recognition of viral dsRNA (38). Based on the comparison of SARS-CoV-2 NSP15 with those of the other 11 coronaviruses, including SARS-CoV and MERS-CoV (38), we speculate that the residue H234 is involved in endoribonuclease activity. Interestingly, the NSP1 but not the NSP15 mutant formed smaller plaques compared to WT SARS-CoV-2 (Fig. 1C) suggesting different effects of the proteins on viral replication.

In COVID-19 patients, SARS-CoV-2 inhibits IFN-I response and reduces the numbers of DC, and both effects contribute to the progression of a severe COVID-19 (99). Additionally, in COVID-19 convalescent patients, the DC maturation markers are suppressed for months after the onset of the disease (55, 100). Strikingly, the NSP1 and NSP15 mutants which demonstrated the greatest IFN-I signaling (Fig. 2) induced the greatest DC maturation (Fig. 3). These effects are consistent with the role of IFN-I in COVID-19 patients, in which a highly impaired IFN-I response was associated with a persistent blood viral load and an exacerbated inflammatory response (10).

Based on the screening of the mutants for their ability to suppress IFN-I response and maturation of DCs, we selected the NSP1 and NSP15 mutants for in-depth characterization *in vitro* and *in vivo*. Infection of Calu-3 cells with the NSP1 and NSP15 mutants activated a set of genes that are linked to immune functions and inflammatory process (Fig. 4), indicating that SARS-CoV-2 interferes with protective immune responses through these proteins. Notably, many genes upregulated by the NSP1 and NSP15 mutant viruses are responsive to IFN-I stimulation (ISG) and are involved in innate defense against viruses, underlying the role of NSP1 and NSP15 in IFN-I antagonism.

Evaluation of the two selected SARS-CoV-2 mutants NSP1 and NSP15 in K18 hACE-2 transgenic mice demonstrated the reduced viral load, inflammation and pathogenicity as compared to the WT virus (Fig. 5). Analysis of the lung transcriptome from the NSP1 mutant-infected mice demonstrated marked changes in pathways associated with regulation of T cell-mediated immunity, inflammatory response, and cytokine production (Fig. 6). The NSP15 mutant gave a more straightforward change: upregulation of defense response, immune effector processes and inflammatory response. These data clearly suggest that NSP1 and NSP15 mutant viruses induced mild to moderate inflammation, as opposed to the severe inflammation noted in mice infected with WT virus. Mice infected with NSP1 and NSP15 mutants had less inflammatory cells in the lungs on histological analysis when compared to the WT virus. The monocytes in the lungs of mice infected with the mutant viruses, especially the NSP15 mutant, had upregulation of several genes and pathways related to immune effector functions (Fig. 7, 8), compared to the WT virus infected animals. Overall, these data indicate that NSP1 and NSP15 inhibit the IFN-I pathways and activation of effector immune cells, contributing to severe disease.

The bulk RNA-seq (Fig. 6) and scRNA-seq (Fig. 7, 8) demonstrated that the NSP1 and NSP15 mutant viruses differentially activated the genes related to the inflammatory process, T cell-mediated immunity, immune effector functions, and positive regulation of cytokine production. scRNA-seq revealed a higher proportion of monocytes and DCs during NSP1 and NSP15 mutant infections, which were associated with reduced lethality (Fig. 5). Importantly, a proportion of inflammatory monocytes was higher in the lungs of mice infected with the NSP1 and NSP15 mutant viruses when compared to those mice infected with WT virus (Fig. 7D), suggesting a protective role of these cells. Multiple gene pathways related to the protective innate immunity, positive regulation of inflammatory response, IFN-I signaling, T cell-mediated immunity, and immune effector functions were activated in mice infected with the NSP1 and NSP15 mutant viruses (Fig. 6, 8). The effects of NSP1 or NSP15 mutations on these responses suggests specific immunosuppressive mechanisms of the two viral proteins. Interestingly, scRNA-seq analysis identified downregulation of NOD pathways in B cells, CD8+ T cells and NK cells after infection with the NSP1 and NSP15 mutants (Fig. 8). This suggests that NSP1 and NSP15 exhibit a control over inflammasome activation. This could also explain the observed reduced pathogenicity of the mutated viruses.

The study has limitations. It did not result in the assessment of the contribution of NSP5 and NSP6 in the viral immunosuppression, as the viruses with the mutations in these genes were not recoverable. Furthermore, immunosuppressive effects of additional SARS-CoV-proteins, which were not included in the study due to the lack of information on their possible immunosuppressive effect when the project was conceived, cannot be excluded. Finally, inflammatory mechanisms involving IL-1 and IL-1ra are different in human and mouse cells (101).

In conclusion, this work demonstrates both distinct and redundant mechanisms of immunosuppression of SARS-CoV-2 mediated by multiple individual viral proteins, with 9 out of the 12 tested proteins showing some immunosuppressive effect in at least one experimental system (Fig. 9). The demonstrated immunosuppressive effects extend from the innate response to immune cells to pathologic changes *in vivo*. Importantly, this work shows, for the first time, a comparison of the effects of multiple viral proteins in the context of authentic viral infection, rather than in a surrogate system, and shows the relative contribution of each viral protein under identical experimental conditions, as opposed to different experimental systems in individual studies. Moreover, information from this study opens the feasibility of testing these mutations, alone or in combination with other attenuating approaches, for the development of live-attenuated vaccines for the prevention of SARS-CoV-2 to effectively control the persisting COVID-19 pandemic.

**Fig. 9.**
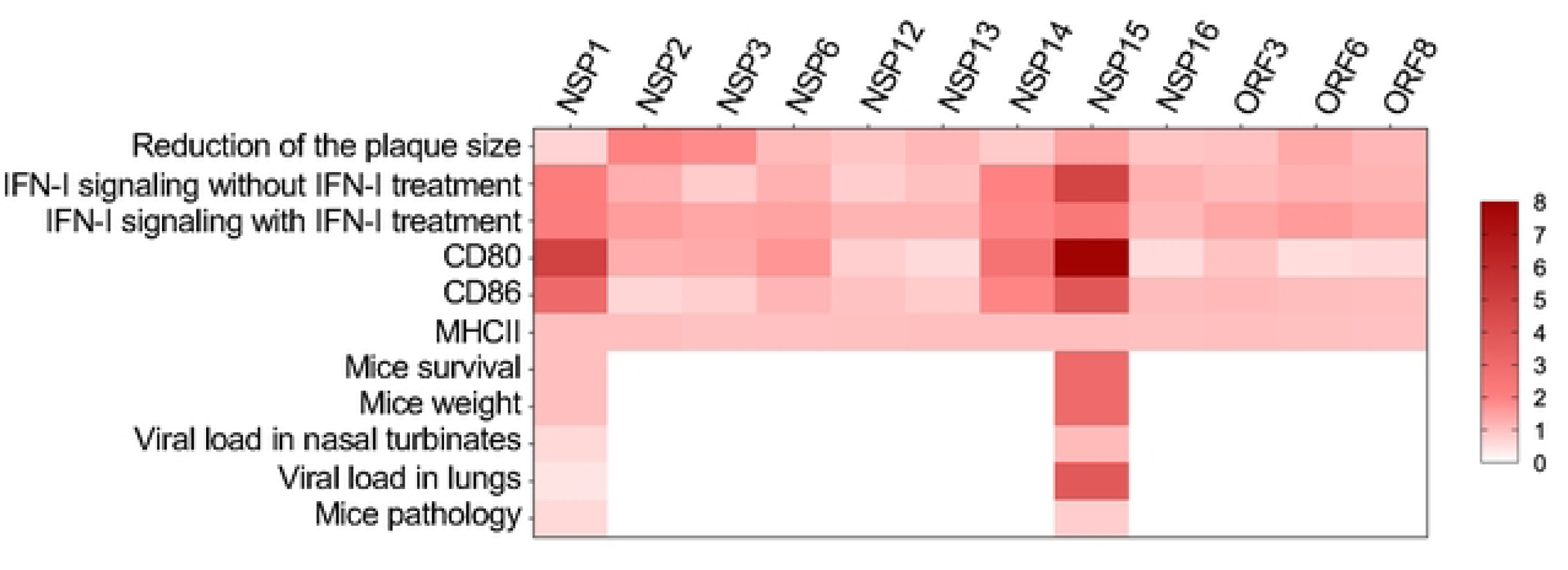
Contribution of individual SARS-CoV-2 proteins in viral immunosuppression and pathogenicity. The figure summarizes the effects of the 12mutated proteins on the viral phenotype andinnate and adaptive immune responses shown in Fig. 1, 2, 3 and 5. The darker colors reflect thegreater effects of the mutations on the biological effects indicated at the left and therefore greater contribution of the corresponding proteins in these biological effects. The transcriptional effects in human cells andin mice are summarized in Fig. 6, 7, 8 and are not included in this heat map.

## METHODS

### Generation of mutated SARS-CoV-2 constructs

The full-length SARS-CoV-2 cDNA clone pBeloBAC11-SARS-CoV-2 (51, 102) was transformed into *E.coli* strain GS1783 (kindly provided by Dr. Gregory A. Smith), and a lambda red system based En Passant mutagenesis was adapted to introduce point mutations or deletions. The Red system, which originates from λ phages, allows for the insertion of linear double-stranded DNA molecules by homologous recombination, and it consists of three proteins named Exo, Beta, and Gam (103, 104). The method uses short identical DNA sequence-mediated recombination between linear DNA and circular DNA to insert DNA sequences into BAC DNA and to remove the marker cassette from the BAC DNA in a second Red recombination step in *e.coli* strain GS1783 with expression of Lambda Red recombination genes (Exo, Beta, and Gam) or/and the I-SceI enzyme under the control of temperature-inducible and arabinose-induced promoters, respectively (52). The final constructs with the selected mutations were generated through the two rounds of recombination. In brief, fragments were amplified from pKan-S with primers, containing the desired mutations, the sequences homologous to I-SceI/Kan resistance cassette and the gene intended to be mutated. The primers used to amplify the I-SceI/Kan^r^ cassette to introduce mutations are shown in Suppl. Table 1. The first recombination between the PCR fragment and pBeloBAC11-SARS-CoV-2 resulted in insertion of the I-SceI/Kan^r^ cassette under kanamycin selection. The cassette carries a I-SceI site and is flanked by two identical short sequences with the desired mutation. The second round of recombination between the two identical sequences removes the I-SceI/Kan^r^ cassette, resulting in the construct with the desired mutations only. To confirm the mutations, the mutated region was PCR-amplified from the mutated constructs for DNA sequencing. To further validate the mutated full-length SARS-CoV-2, DNA was characterized, in parallel with the parental non-mutated clone, by BsmBI restriction endonuclease.

### Recovery of SARS-CoV-2 mutants

The full-length clone pBeloBAC11-SARS-CoV-2 and its mutated derivatives were used for recovery of SARS-CoV-2 mutants using the previously described method (51, 102) with some modifications. The procedure was performed under BSL-3 containment of the Galveston National Laboratory. Briefly, 80% confluent monolayers of Vero E6 cells in 12-well plates were transfected with 1.0 μg per well of infectious SARS-CoV-2 BAC DNA or its mutated derivatives using TransIT®-LT1 transfection reagent according to manufacturer’s instructions (Mirus Bio). At 48 h post transfection, the transfected cells were split and seeded into 25 cm^2^ flasks with 50% confluent monolayers of Vero E6 cells. The cell cultures were observed daily until the appearance of a strong cytopathic effect to collect viral supernatants. To confirm the desired mutation in the recombinant viral mutants, DNA-free viral RNA was isolated from the viral supernatants using Direct-Zol RNA Microprep kit (Zymo research) according to manufacturer’s instruction. The mutations in the viral genomes were further confirmed by reverse transcription-PCR followed by Sanger DNA sequencing.

### Virus titration

Vero E6 cells were seeded on 96-well plates or 12-well plates. On the next day, the cells were incubated with 10-fold serial dilutions of the virus for 1 h and overlaid with 0.55% methylcellulose or 1.0% cellulose in minimal essential medium (MEM) (Gibco) supplemented with 2% FBS (Corning) and 50 µg/ml gentamycin (Corning/Fisher). If methylcellulose was used, the overlay was discarded, the infected cells were fixed with 10% formalin for 24 h, then stained with 10% formalin containing 0.25% crystal violet at room temperature for 15 min and washed with water or stained with a mixture of 37 human monoclonal antibodies targeting different antigenic sites within spike protein of SARS-CoV-2 (105). Then the plaques were visualized using HRP-conjugated anti-human IgG with Nova-red substrate kit (Vector Laboratories). If cellulose was used, the overlay was discarded, and the infected cells were fixed with 10% formalin for 24 h, and the monolayers were stained with 10% formalin containing 0.25% crystal violet at room temperature for 15 min and washed with water.

### ISRE-luciferase reporter assay

293T-ACE2/TMPRSS2 cells were plated onto poly-L-lysin-coated 96 well plates (Corning, cat. #3917), and one day later the cells were transfected with pISRE-luc and pTK-RL (Promega) using TransIT®-LT1 Transfection Reagent. pISRE-luc has a firefly luciferase gene driven by a minimal promoter with IFN-stimulated responsive element (ISRE) and pTK-R encodes a constitutively expressed *Renilla* luciferase as a transfection control. Next day, the cells were mock infected or infected with SARS-CoV-2 or its mutants at 0.3 PFU/cell. At 1 day after the infection the cells were untreated or treated with universal IFN (PBL Assay Science) at 100 uints/ml. At 1 day after the treatment the reporter assay was performed using Dual-Glo Luciferase System (Promega, Cat# E2920) according to manufacturer’s instructions. The luminescence in the wells of the 96-well plates was measured using the Cytation 7 reader (BioTek Instruments).

### Human dendritic cell maturation assay

Human peripheral blood mononuclear cells (PBMC) were used for isolation of primary monocytes. The PBMC were collected by gradient centrifugation using Histopaque-1077 (Sigma) from the commercially sourced buffy coats (Gulf Coast Regional Blood Center, Houston, Texas, USA) of three healthy donors. The monocytes were isolated from PBMCs by magnetic positive selection with MidiMacs separation columns (Miltenyi Biotech) and anti-CD14 antibody coated magnetic beads (Miltenyi Biotech). Purified monocytes were cultured in monocyte-derived DC differentiation medium containing IL-4 and GM-CSF (Miltenyi Biotech) for 6 days at 37°C in humidified air containing 5% CO_2_. Cells were replenished with fresh medium on day 3. The procedure resulted in differentiation of monocytes into monocyte-derived DCs with an immature phenotype (CD11c^+^/CD14^-^). On day 6, DCs were harvested using enzyme-free PBS-based cell dissociation buffer (Gibco) and washed in RPMI-1640 medium (Gibco) supplemented with 10% FCS, 1% penicillin/streptomycin and 1% pyruvate. Cells were counted and aliquoted in duplicates in 5 ml Falcon tubes (5 x 10^4^ cells in 100 µl medium per tube). Cells were treated with LPS (1 ng/ml) or SARS-CoV-2 (MOI 3 PFU/cell) under BSL-3 containment and cultured for 1, 2 or 3 days. After incubation, DC maturation status was analyzed by multicolor flow cytometry using anti-human antibodies specific for for CD11c (clone BU15), CD80 (clone W17149D), CD86 (clone BU63) and MHCII/HLA-DR (clone LN3) (Biolegend). Cells were incubated with a cocktail of antibodies for 10 min at room temperature, washed once with PBS, fixed twice with 4% PFA and removed from BSL3 containment. The fixed cells were analyzed using the LSRFortessa flow cytometer (BD Biosciences). After exclusion of cell debris from the analysis by forward and side scatter gating, the cells were gated for CD11c+ population which were further gated for the expression of CD80, CD86 or HLA-DR. The percentages of cells expressing the maturation markers and their MFI values were calculated in FlowJo software v10.8 (BD).

### Isolation of RNA from infected Calu-3 cells or homogenized mouse lungs

The infected cells or homogenized lungs were lysed in Trizol to prepare total RNAs using Direct-Zol RNA Microprep kit (Zymo Research) with DNase treatment according to the manufacturer’s instructions. The isolated RNA was submitted for RNA sequencing.

### Preparation of cDNA library and analysis of the RNA-seq data

For Calu-3 cell-derived RNA, the concentration and integrity (RIN) of isolated RNA were determined using the Quant-iT™ RiboGreen™ RNA Assay Kit (Thermo Fisher) and an RNA Standard Sensitivity Kit (Agilent Technologies, catalog number DNF-471) on a Fragment Analyzer Automated CE system (Agilent Technologies). Subsequently, cDNA libraries were constructed from total RNA using the Universal Plus mRNA-Seq kit (Tecan Genomics) in a Biomek i7 Automated Workstation (Beckman Coulter). Briefly, mRNA was isolated from purified 300 ng total RNA using oligo-dT beads and used to synthesize cDNA following the manufacturer’s instructions. The transcripts for ribosomal RNA (rRNA) and globin were further depleted using the AnyDeplete kit (Tecan Genomics) prior to the amplification of libraries. Library concentration was assessed fluorometrically using the Qubit dsDNA HS Kit (Thermo Fisher), and quality was assessed with the HS NGS Fragment Kit (1–6,000bp; DNF-474, Agilent Technologies).

Following library preparation, samples were pooled and preliminary sequencing of cDNA libraries (average read depth of 90,000 reads) was performed using a MiSeq system (Illumina), to confirm library quality and concentration. Deep sequencing was subsequently performed using an S4 flow cell in a NovaSeq sequencing system (Illumina; average read depth ∼30 million pairs of 2×100bp reads) at the New York Genome Center. Reads were subjected to quality control using FastQC (v0.11.8) and RNASeqMetrics (Picard v2.18.16), mapped using STAR (v2.7.0d), and gene counts were summarized using RSEM (v1.3.1). Counts were normalized using DESeq2 (v1.30.1) regularized-log transform function, following a differential gene expression analysis. Gene Ontology enrichment analyses were performed using the R packages Go.db (v.3.7.0) and GOstats (2.48.0).

Lung derived mRNA sequencing was conducted according to the published SMART-3SEQ method (106, 107). Concisely, the first strand primer was annealed to 1 µl of sample RNA before being extended with SMARTScribe reverse transcriptase (Clontech). After the addition of the second strand primer, second strand synthesis was carried out, and adapter sequences with unique indexes were inserted with 15 cycles of PCR using NEB Next single index adapters (New England BioLabs). Purified PCR products were measured and pooled for sequencing on an Illumina NextSeq 550 High Output Flow Cell using the single-end 75 base technique using AMPure XP SPRI beads (Beckman Coulter Life Sciences).

A five-base unique molecular identification (UMI) and 3Gs were added to the 5′ end of every sequence using the Smart-3SEQ protocol. Using STAR version 2.7.5c and the settings suggested by the software developers for the Encode consortium, reads were aligned to the Mus musculus NCBI assembly GCF_000349665.1. The NCBI annotation version 102 was utilized to count reads per gene using the FeatureCounts software. The count table was utilized as an input into DESeq2, allowed for the estimation of differential gene expression in accordance with the DESeq2 vignette that were provided with the software. The R heatmap tool was used to do hierarchical clustering of the genes. Further, Metascape was used to perform pathway enrichment analysis on the set of differentially expressed genes with p values <0.05. The enriched pathways were identified using the Gene Ontology (GO) terms biological process, molecular function, and KEGG pathway. Pathways were chosen with an FDR adjusted p value < 0.01. All heatmaps were created using the Python 3.9 libraries matplotlib and Seaborn.

### Infection of K18 hACE2 transgenic mice

The mouse experiments were approved by the UTMB Institutional Care and Use Committee (IACUC) and the Texas Biomedical Research Institute IACUC and conducted in an animal biosafety level 3 (ABSL-3) facility of the Galveston National laboratory and the Texas Biomedical Research Institute. The K18 hACE2 transgenic mice B6.Cg-Tg(K18-ACE2)2Prlmn/J were purchased from the Jackson Laboratory. The following three experiments were conducted:

1. Infection of mice to assess the survival and disease. Six-week-old K18 hACE2 transgenic mice at 4 animals per group, total of 16 mice, were sedated with isoflurane and either mock-infected with medium or infected intranasally with 10^5^ PFU of SARS-CoV-2 or the mutants with a final volume of 50 µl (∼25 μl into each nostril). After infection, mice were monitored daily for survival and body weight. Mice showing >25% loss of their initial body weight were defined as reaching the experimental endpoint and humanely euthanized.
2. Infection of mice to assess viral load and change in gene expression in lungs. Six-week-old female K18 hACE2 transgenic mice at 8 animals per group, total of 32 mice, were either mock-infected with medium or infected as described above. The mice were euthanized at 2 and 4 days post-infection, and the lungs were isolated for assessment of gross changes. The whole lungs were excised and photographed. Next, the left lungs were fixed in 10% neutral buffered formalin for histopathology, and the right lungs were processed for virus titration and RNA isolation as follows. The right lung was homogenized in Precellys tubes, then frozen in a -80°C freezer. After thawing the tubes were centrifuged at 12,000 x g at 4°C for 5 min, and supernatants were collected for virus titration. The pellets were lysed in 1 ml Trizol for RNA isolation according to the manufacturer’s instructions.
3. Infection of mice for single cell RNA sequencing of lung immune cells. Five-to seven-week-old K18 hACE2 transgenic mice (4 mice per group, total of 16 mice) were either mock-infected with medium or infected as described above. On day 3 mice were euthanized as described above, and the lungs were isolated for immune cell isolation.

### Single cell isolation and RNA sequencing

Single-cell isolation and RNA sequencing were performed as described previously (108). In brief, the freshly collected whole lungs were dissected into single lobes to isolate single cells using a mouse lung dissociation kit (Miltenyi Biotec) and a gentleMACS™ Dissociator (Miltenyi Biotec), and red blood cells were lysed with 1X RBC lysis buffer (eBioscience). Cell suspensions were processed for single-cell sequencing following the protocol for Chromium Next GEM Single Cell 5’ Version 2 (dual index). 10,000 cells were targeted. The transcriptome of each cell was indexed with a pool of 750,000 barcodes by partitioning each cell into Gel beads-in-emulsion (GEMs) combined with a Master Mix containing reverse transcription (RT) reagents and poly(dT) RT primers. The emulsion was generated using Next GEM chips and the Chromium Controller device (10x Genomics). The RT reaction to the generated emulsion produced 10x Barcoded full-length cDNA from poly-adenylated mRNA. This initial cDNA was PCR amplified to produce material for 5’ gene expression sequencing. After PCR amplification, bioanalyzer Quality Control (QC) was performed for all the samples using the Agilent Bioanalyzer High Sensitivity DNA assay in the 2100 expert software (Agilent). All the samples passed the initial QC with a cDNA size 700-1,500 bp.

Amplified full-length cDNA from polyadenylated mRNA was used to generate 5’ gene expression (GEX) libraries. The cDNA was enzymatically fragmented, and size selected to optimize the cDNA amplicon size. P5, P7, i5 and i7 sample indexes, and Illumina R2 sequences (read 2 primer sequences) were added via End Repair, A-tailing, Adaptor ligation and sample index PCR. A second QC was performed for each library before sequencing with an expected library size 500 – 900 bp. Finally, libraries were pooled and sequenced by the New York Genome Center (NYGC) using a NovaSeq sequencer and S2 flowcell with a minimum of 20,000 reads pair per cell.

### scRNA sequencing data processing

To enable quantification of both viral and host genes from single cell data, we first merged the mm10(GRCm38.p6) and SARS-CoV-2 reference genomes (ASM993790v1). Cell Ranger v7.0.0 (10× Genomics) was then used to demultiplex cellular barcodes and align the reads to the combined genome. The filtered feature-barcode matrix per sample was then read into a Seurat object (109). Putative doublets from each sample were identified using scds (110). Any cell with the scds hybrid doublet score of 0.8 (doublet scoring based on co-expression and binary classification) was considered a putative doublet and was removed. Additionally, cells with unusually high expression levels (nFeature over 4,000 or nCount over 10,000) were also considered putative doublets and were removed from further analysis. Poor quality cells (nFeature of less than 1,000 or over 5% mitochondrial content) were also filtered out. The merged Seurat dataset was examined for the presence of intrinsic batches. All the samples were then integrated following the Seurat integration pipeline while correcting for the identified batch labels.

Initial cell-type annotation was performed with SingleR using the reference from the Immunologic Genome Project, accessed via celldex (111). Cell cycle (G2M and S phases) scores were calculated using the Seurat function CellCycleScoring. A single cluster with high G2M and S phase scores was identified and labeled as ‘Proliferating’. Putative cell type annotation of each cluster was also validated using canonical marker expression (Fig. 7D and Suppl. Fig. 5). Major clusters corresponding to 15 cell types were identified among single cells which were isolated from lung tissue. The average expression of SARS-CoV-2 viral genes in each cell type/sample pair were also calculated (Suppl. Fig. 8).

Within each cell type, differential expression analysis was performed between the following sample groups: WT vs mock, NSP1 mutant vs WT, and NSP15 mutant vs WT. Differentially expressed genes (DEG) at single cell level were identified using the Wilcoxon test. Those passing the FDR-adjusted p-value < 0.05 and absolute log2 fold change > 0.3 in expression were selected as significant. We also performed a DEG analysis at the pseudo-bulk per cell-type level. Because of the small sample sizes, this analysis did not have sufficient power to detect differentially expressed genes and was not used further.

Pathway enrichment analysis was performed on the sets of DEG with enrichR (112) using the Gene Ontology (GO) biological process terms. All terms with FDR adjusted p-value < 0.1 were considered significant. Since GO is a hierarchy with closely related terms that correspond to highly overlapping gene sets, we sub-selected significant terms for presentation in Fig. 8A, B and Suppl. Fig. 10. The sub-selection was done as follows: terms were included if there was a difference of at least two regulated genes from every other term. The direction of regulation for each enriched term was determined by the proportion of upregulated DEG versus the downregulated DEG for each cell type.

### Integration of the non-genomic data in the heat map

Heatmap depicting the overall changes in different parameters among all SARS-CoV2 mutants. Mean of individual values of Plaque size, IFN response without treatment, IFN response with treatment, DC CD80, DC CD86 and DC MHC-II was calculated for each mutant followed by normalization with the mean WT SARS-CoV-2 control. For further investigation, several other parameters including mice survival, mice weight, viral load in NL, viral load in lungs and mice pathology were studied in NSP1 and NSP15 mutant strain only. Mean of individual values of the mentioned parameters was calculated for both mutants followed by normalization with the mean of WT SARS-CoV-2. Heatmap was generated through GraphPad prism (version 9.5.1).

## ACKNOWLEDGEMENTS

We thank the UTMB Animal Resource Center veterinary staff for the technical support of mouse experiments. We thank Dr. Gregory A. Smith at Northwestern University for providing *E.coli* strain GS1783 and Dr. Mohsan Saeed at Boston University for providing 293T-ACE2/TMPRSS2 cells. We thank Dr. Natalia Kuzmina for help with isolation of immune cells from mouse lungs. This project was partially funded by UTMB intramural funds and partially by Defense Advanced Research Project Agency (DARPA) Grant N6600119C4022 (to S.C.S and A.B.).

## AUTHOR RESEARCH CONTRIBUTIONS

Conceptualizaiton: F.Z., S.C.S, and A.B. Methodology: F.Z., S.P., N.J., W.S.C., R.S.A., P.A.I., C.Y., N.A.K., S.C., G.N., E.Z., S.G.W., L.M.-S., S.C.S, and A.B. Formal analysis: F.Z., S.P., W.S.C, R.S.A., P.A.I., S.C., G.N., and S.G.W. Investigation: all authors. Writing – original draft: F.Z., S.P., W.S.C., S.C., and A.B. Writing – review and editing: all authors. Supervision: A.B., L.M.-S. and S.C.S. Project administration: A.B. Funding acquisition: A.B., L.M.-S. and S.C.S.

## Notes

### Competing Interest Statement

The authors have declared no competing interest.

## REFERENCES

1. WHO 2020, posting date. Coronavirus disease (COVID-19) Pandemic. https://covid19.who.int.

2. Harvey WT, Carabelli AM, Jackson B, Gupta RK, Thomson EC, Harrison EM, Ludden C, Reeve R, Rambaut A, Consortium C-GU, Peacock SJ, Robertson DL. 2021. SARS-CoV-2 variants, spike mutations and immune escape. Nat Rev Microbiol 19:409–424.

3. Hamming I, Timens W, Bulthuis ML, Lely AT, Navis G, van Goor H. 2004. Tissue distribution of ACE2 protein, the functional receptor for SARS coronavirus. A first step in understanding SARS pathogenesis. J Pathol 203:631–637.

4. Lamers MM, Haagmans BL. 2022. SARS-CoV-2 pathogenesis. Nat Rev Microbiol 20:270–284.

5. Moustaqil M, Ollivier E, Chiu HP, Van Tol S, Rudolffi-Soto P, Stevens C, Bhumkar A, Hunter DJB, Freiberg AN, Jacques D, Lee B, Sierecki E, Gambin Y. 2021. SARS-CoV-2 proteases PLpro and 3CLpro cleave IRF3 and critical modulators of inflammatory pathways (NLRP12 and TAB1): implications for disease presentation across species. Emerg Microbes Infect 10:178–195.

6. V’Kovski P, Kratzel A, Steiner S, Stalder H, Thiel V. 2021. Coronavirus biology and replication: implications for SARS-CoV-2. Nat Rev Microbiol 19:155–170.

7. Khorramdelazad H, Kazemi MH, Azimi M, Aghamajidi A, Mehrabadi AZ, Shahba F, Aghamohammadi N, Falak R, Faraji F, Jafari R. 2022. Type-I interferons in the immunopathogenesis and treatment of Coronavirus disease 2019. Eur J Pharmacol 927:175051.

8. Masood KI, Yameen M, Ashraf J, Shahid S, Mahmood SF, Nasir A, Nasir N, Jamil B, Ghanchi NK, Khanum I, Razzak SA, Kanji A, Hussain R, M ER, Hasan Z. 2021. Upregulated type I interferon responses in asymptomatic COVID-19 infection are associated with improved clinical outcome. Sci Rep 11:22958.

9. Ziegler CGK, Miao VN, Owings AH, Navia AW, Tang Y, Bromley JD, Lotfy P, Sloan M, Laird H, Williams HB, George M, Drake RS, Christian T, Parker A, Sindel CB, Burger MW, Pride Y, Hasan M, Abraham GE, 3rd, Senitko M, Robinson TO, Shalek AK, Glover SC, Horwitz BH, Ordovas-Montanes J. 2021. Impaired local intrinsic immunity to SARS-CoV-2 infection in severe COVID-19. Cell 184:4713–4733 e4722.

10. Hadjadj J, Yatim N, Barnabei L, Corneau A, Boussier J, Smith N, Pere H, Charbit B, Bondet V, Chenevier-Gobeaux C, Breillat P, Carlier N, Gauzit R, Morbieu C, Pene F, Marin N, Roche N, Szwebel TA, Merkling SH, Treluyer JM, Veyer D, Mouthon L, Blanc C, Tharaux PL, Rozenberg F, Fischer A, Duffy D, Rieux-Laucat F, Kerneis S, Terrier B. 2020. Impaired type I interferon activity and inflammatory responses in severe COVID-19 patients. Science 369:718–724.

11. Lopez J, Mommert M, Mouton W, Pizzorno A, Brengel-Pesce K, Mezidi M, Villard M, Lina B, Richard JC, Fassier JB, Cheynet V, Padey B, Duliere V, Julien T, Paul S, Bastard P, Belot A, Bal A, Casanova JL, Rosa-Calatrava M, Morfin F, Walzer T, Trouillet-Assant S. 2021. Early nasal type I IFN immunity against SARS-CoV-2 is compromised in patients with autoantibodies against type I IFNs. J Exp Med 218.

12. Cheemarla NR, Watkins TA, Mihaylova VT, Wang B, Zhao D, Wang G, Landry ML, Foxman EF. 2021. Dynamic innate immune response determines susceptibility to SARS-CoV-2 infection and early replication kinetics. J Exp Med 218.

13. Beyer DK, Forero A. 2022. Mechanisms of Antiviral Immune Evasion of SARS-CoV-2. J Mol Biol 434:167265.

14. Thoms M, Buschauer R, Ameismeier M, Koepke L, Denk T, Hirschenberger M, Kratzat H, Hayn M, Mackens-Kiani T, Cheng J, Straub JH, Sturzel CM, Frohlich T, Berninghausen O, Becker T, Kirchhoff F, Sparrer KMJ, Beckmann R. 2020. Structural basis for translational shutdown and immune evasion by the Nsp1 protein of SARS-CoV-2. Science 369:1249–1255.

15. Xia H, Cao Z, Xie X, Zhang X, Chen JY, Wang H, Menachery VD, Rajsbaum R, Shi PY. 2020. Evasion of Type I Interferon by SARS-CoV-2. Cell Rep 33:108234.

16. Banerjee AK, Blanco MR, Bruce EA, Honson DD, Chen LM, Chow A, Bhat P, Ollikainen N, Quinodoz SA, Loney C, Thai J, Miller ZD, Lin AE, Schmidt MM, Stewart DG, Goldfarb D, De Lorenzo G, Rihn SJ, Voorhees RM, Botten JW, Majumdar D, Guttman M. 2020. SARS-CoV-2 Disrupts Splicing, Translation, and Protein Trafficking to Suppress Host Defenses. Cell 183:1325–1339 e1321.

17. Miorin L, Kehrer T, Sanchez-Aparicio MT, Zhang K, Cohen P, Patel RS, Cupic A, Makio T, Mei M, Moreno E, Danziger O, White KM, Rathnasinghe R, Uccellini M, Gao S, Aydillo T, Mena I, Yin X, Martin-Sancho L, Krogan NJ, Chanda SK, Schotsaert M, Wozniak RW, Ren Y, Rosenberg BR, Fontoura BMA, Garcia-Sastre A. 2020. SARS-CoV-2 Orf6 hijacks Nup98 to block STAT nuclear import and antagonize interferon signaling. Proc Natl Acad Sci U S A 117:28344–28354.

18. Shemesh M, Aktepe TE, Deerain JM, McAuley JL, Audsley MD, David CT, Purcell DFJ, Urin V, Hartmann R, Moseley GW, Mackenzie JM, Schreiber G, Harari D. 2021. SARS-CoV-2 suppresses IFNbeta production mediated by NSP1, 5, 6, 15, ORF6 and ORF7b but does not suppress the effects of added interferon. PLoS Pathog 17:e1009800.

19. Lin JW, Tang C, Wei HC, Du B, Chen C, Wang M, Zhou Y, Yu MX, Cheng L, Kuivanen S, Ogando NS, Levanov L, Zhao Y, Li CL, Zhou R, Li Z, Zhang Y, Sun K, Wang C, Chen L, Xiao X, Zheng X, Chen SS, Zhou Z, Yang R, Zhang D, Xu M, Song J, Wang D, Li Y, Lei S, Zeng W, Yang Q, He P, Zhang Y, Zhou L, Cao L, Luo F, Liu H, Wang L, Ye F, Zhang M, Li M, Fan W, Li X, Li K, Ke B, Xu J, Yang H, He S, Pan M, Yan Y, Zha Y, Jiang L, Yu C, Liu Y, Xu Z, Li Q, Jiang Y, Sun J, Hong W, Wei H, Lu G, Vapalahti O, Luo Y, Wei Y, Connor T, Tan W, Snijder EJ, Smura T, Li W, Geng J, Ying B, Chen L. 2021. Genomic monitoring of SARS-CoV-2 uncovers an Nsp1 deletion variant that modulates type I interferon response. Cell Host Microbe 29:489–502 e488.

20. Konno Y, Kimura I, Uriu K, Fukushi M, Irie T, Koyanagi Y, Sauter D, Gifford RJ, Consortium U-C, Nakagawa S, Sato K. 2020. SARS-CoV-2 ORF3b Is a Potent Interferon Antagonist Whose Activity Is Increased by a Naturally Occurring Elongation Variant. Cell Rep 32:108185.

21. Kumar A, Ishida R, Strilets T, Cole J, Lopez-Orozco J, Fayad N, Felix-Lopez A, Elaish M, Evseev D, Magor KE, Mahal LK, Nagata LP, Evans DH, Hobman TC. 2021. SARS-CoV-2 Nonstructural Protein 1 Inhibits the Interferon Response by Causing Depletion of Key Host Signaling Factors. J Virol 95:e0026621.

22. Li JY, Liao CH, Wang Q, Tan YJ, Luo R, Qiu Y, Ge XY. 2020. The ORF6, ORF8 and nucleocapsid proteins of SARS-CoV-2 inhibit type I interferon signaling pathway. Virus Res 286:198074.

23. Lei X, Dong X, Ma R, Wang W, Xiao X, Tian Z, Wang C, Wang Y, Li L, Ren L, Guo F, Zhao Z, Zhou Z, Xiang Z, Wang J. 2020. Activation and evasion of type I interferon responses by SARS-CoV-2. Nat Commun 11:3810.

24. Jiang HW, Zhang HN, Meng QF, Xie J, Li Y, Chen H, Zheng YX, Wang XN, Qi H, Zhang J, Wang PH, Han ZG, Tao SC. 2020. SARS-CoV-2 Orf9b suppresses type I interferon responses by targeting TOM70. Cell Mol Immunol 17:998–1000.

25. Yuen CK, Lam JY, Wong WM, Mak LF, Wang X, Chu H, Cai JP, Jin DY, To KK, Chan JF, Yuen KY, Kok KH. 2020. SARS-CoV-2 nsp13, nsp14, nsp15 and orf6 function as potent interferon antagonists. Emerg Microbes Infect 9:1418–1428.

26. Mu J, Fang Y, Yang Q, Shu T, Wang A, Huang M, Jin L, Deng F, Qiu Y, Zhou X. 2020. SARS-CoV-2 N protein antagonizes type I interferon signaling by suppressing phosphorylation and nuclear translocation of STAT1 and STAT2. Cell Discov 6:65.

27. Gordon DE, Jang GM, Bouhaddou M, Xu J, Obernier K, White KM, O’Meara MJ, Rezelj VV, Guo JZ, Swaney DL, Tummino TA, Huttenhain R, Kaake RM, Richards AL, Tutuncuoglu B, Foussard H, Batra J, Haas K, Modak M, Kim M, Haas P, Polacco BJ, Braberg H, Fabius JM, Eckhardt M, Soucheray M, Bennett MJ, Cakir M, McGregor MJ, Li Q, Meyer B, Roesch F, Vallet T, Mac Kain A, Miorin L, Moreno E, Naing ZZC, Zhou Y, Peng S, Shi Y, Zhang Z, Shen W, Kirby IT, Melnyk JE, Chorba JS, Lou K, Dai SA, Barrio-Hernandez I, Memon D, Hernandez-Armenta C, Lyu J, Mathy CJP, Perica T, Pilla KB, Ganesan SJ, Saltzberg DJ, Rakesh R, Liu X, Rosenthal SB, Calviello L, Venkataramanan S, Liboy-Lugo J, Lin Y, Huang XP, Liu Y, Wankowicz SA, Bohn M, Safari M, Ugur FS, Koh C, Savar NS, Tran QD, Shengjuler D, Fletcher SJ, O’Neal MC, Cai Y, Chang JCJ, Broadhurst DJ, Klippsten S, Sharp PP, Wenzell NA, Kuzuoglu-Ozturk D, Wang HY, Trenker R, Young JM, Cavero DA, Hiatt J, Roth TL, Rathore U, Subramanian A, Noack J, Hubert M, Stroud RM, Frankel AD, Rosenberg OS, Verba KA, Agard DA, Ott M, Emerman M, Jura N, von Zastrow M, Verdin E, Ashworth A, Schwartz O, d’Enfert C, Mukherjee S, Jacobson M, Malik HS, Fujimori DG, Ideker T, Craik CS, Floor SN, Fraser JS, Gross JD, Sali A, Roth BL, Ruggero D, Taunton J, Kortemme T, Beltrao P, Vignuzzi M, Garcia-Sastre A, Shokat KM, Shoichet BK, Krogan NJ. 2020. A SARS-CoV-2 protein interaction map reveals targets for drug repurposing. Nature 583:459–468.

28. Xu Z, Choi JH, Dai DL, Luo J, Ladak RJ, Li Q, Wang Y, Zhang C, Wiebe S, Liu ACH, Ran X, Yang J, Naeli P, Garzia A, Zhou L, Mahmood N, Deng Q, Elaish M, Lin R, Mahal LK, Hobman TC, Pelletier J, Alain T, Vidal SM, Duchaine T, Mazhab-Jafari MT, Mao X, Jafarnejad SM, Sonenberg N. 2022. SARS-CoV-2 impairs interferon production via NSP2-induced repression of mRNA translation. Proc Natl Acad Sci U S A 119:e2204539119.

29. Wang W, Zhou Z, Xiao X, Tian Z, Dong X, Wang C, Li L, Ren L, Lei X, Xiang Z, Wang J. 2021. SARS-CoV-2 nsp12 attenuates type I interferon production by inhibiting IRF3 nuclear translocation. Cell Mol Immunol 18:945–953.

30. Russ A, Wittmann S, Tsukamoto Y, Herrmann A, Deutschmann J, Lagisquet J, Ensser A, Kato H, Gramberg T. 2022. Nsp16 shields SARS-CoV-2 from efficient MDA5 sensing and IFIT1-mediated restriction. EMBO Rep 23:e55648.

31. Rashid F, Dzakah EE, Wang H, Tang S. 2021. The ORF8 protein of SARS-CoV-2 induced endoplasmic reticulum stress and mediated immune evasion by antagonizing production of interferon beta. Virus Res 296:198350.

32. Bello-Perez M, Hurtado-Tamayo J, Mykytyn AZ, Lamers MM, Requena-Platek R, Schipper D, Munoz-Santos D, Ripoll-Gomez J, Esteban A, Sanchez-Cordon PJ, Enjuanes L, Haagmans BL, Sola I. 2023. SARS-CoV-2 ORF8 accessory protein is a virulence factor. mBio:e0045123.

33. Zhang K, Miorin L, Makio T, Dehghan I, Gao S, Xie Y, Zhong H, Esparza M, Kehrer T, Kumar A, Hobman TC, Ptak C, Gao B, Minna JD, Chen Z, Garcia-Sastre A, Ren Y, Wozniak RW, Fontoura BMA. 2021. Nsp1 protein of SARS-CoV-2 disrupts the mRNA export machinery to inhibit host gene expression. Sci Adv 7.

34. Schubert K, Karousis ED, Jomaa A, Scaiola A, Echeverria B, Gurzeler LA, Leibundgut M, Thiel V, Muhlemann O, Ban N. 2020. SARS-CoV-2 Nsp1 binds the ribosomal mRNA channel to inhibit translation. Nat Struct Mol Biol 27:959–966.

35. Sui C, Xiao T, Zhang S, Zeng H, Zheng Y, Liu B, Xu G, Gao C, Zhang Z. 2022. SARS-CoV-2 NSP13 Inhibits Type I IFN Production by Degradation of TBK1 via p62-Dependent Selective Autophagy. J Immunol 208:753–761.

36. Feng K, Zhang HJ, Min YQ, Zhou M, Deng F, Wang HL, Li PQ, Ning YJ. 2023. SARS-CoV-2 NSP13 interacts with host IRF3, blocking antiviral immune responses. J Med Virol 95:e28881.

37. Deng X, Baker SC. 2018. An "Old" protein with a new story: Coronavirus endoribonuclease is important for evading host antiviral defenses. Virology 517:157–163.

38. Deng X, Hackbart M, Mettelman RC, O’Brien A, Mielech AM, Yi G, Kao CC, Baker SC. 2017. Coronavirus nonstructural protein 15 mediates evasion of dsRNA sensors and limits apoptosis in macrophages. Proc Natl Acad Sci U S A 114:E4251–E4260.

39. Bouvet M, Debarnot C, Imbert I, Selisko B, Snijder EJ, Canard B, Decroly E. 2010. In vitro reconstitution of SARS-coronavirus mRNA cap methylation. PLoS Pathog 6:e1000863.

40. Xia H, Shi PY. 2020. Antagonism of Type I Interferon by Severe Acute Respiratory Syndrome Coronavirus 2. J Interferon Cytokine Res 40:543–548.

41. Zhang J, Ejikemeuwa A, Gerzanich V, Nasr M, Tang Q, Simard JM, Zhao RY. 2022. Understanding the Role of SARS-CoV-2 ORF3a in Viral Pathogenesis and COVID-19. Front Microbiol 13:854567.

42. Schroeder S, Pott F, Niemeyer D, Veith T, Richter A, Muth D, Goffinet C, Muller MA, Drosten C. 2021. Interferon antagonism by SARS-CoV-2: a functional study using reverse genetics. Lancet Microbe 2:e210–e218.

43. Kopecky-Bromberg SA, Martinez-Sobrido L, Frieman M, Baric RA, Palese P. 2007. Severe acute respiratory syndrome coronavirus open reading frame (ORF) 3b, ORF 6, and nucleocapsid proteins function as interferon antagonists. J Virol 81:548–557.

44. Frieman M, Yount B, Heise M, Kopecky-Bromberg SA, Palese P, Baric RS. 2007. Severe acute respiratory syndrome coronavirus ORF6 antagonizes STAT1 function by sequestering nuclear import factors on the rough endoplasmic reticulum/Golgi membrane. J Virol 81:9812–9824.

45. Hall R, Guedan A, Yap MW, Young GR, Harvey R, Stoye JP, Bishop KN. 2022. SARS-CoV-2 ORF6 disrupts innate immune signalling by inhibiting cellular mRNA export. PLoS Pathog 18:e1010349.

46. Vinjamuri S, Li L, Bouvier M. 2022. SARS-CoV-2 ORF8: One protein, seemingly one structure, and many functions. Front Immunol 13:1035559.

47. Arshad N, Laurent-Rolle M, Ahmed WS, Hsu JC, Mitchell SM, Pawlak J, Sengupta D, Biswas KH, Cresswell P. 2023. SARS-CoV-2 accessory proteins ORF7a and ORF3a use distinct mechanisms to down-regulate MHC-I surface expression. Proc Natl Acad Sci U S A 120:e2208525120.

48. Shin D, Mukherjee R, Grewe D, Bojkova D, Baek K, Bhattacharya A, Schulz L, Widera M, Mehdipour AR, Tascher G, Geurink PP, Wilhelm A, van der Heden van Noort GJ, Ovaa H, Muller S, Knobeloch KP, Rajalingam K, Schulman BA, Cinatl J, Hummer G, Ciesek S, Dikic I. 2020. Papain-like protease regulates SARS-CoV-2 viral spread and innate immunity. Nature 587:657–662.

49. Hayn M, Hirschenberger M, Koepke L, Nchioua R, Straub JH, Klute S, Hunszinger V, Zech F, Prelli Bozzo C, Aftab W, Christensen MH, Conzelmann C, Muller JA, Srinivasachar Badarinarayan S, Sturzel CM, Forne I, Stenger S, Conzelmann KK, Munch J, Schmidt FI, Sauter D, Imhof A, Kirchhoff F, Sparrer KMJ. 2021. Systematic functional analysis of SARS-CoV-2 proteins uncovers viral innate immune antagonists and remaining vulnerabilities. Cell Rep 35:109126.

50. Liu Y, Qin C, Rao Y, Ngo C, Feng JJ, Zhao J, Zhang S, Wang TY, Carriere J, Savas AC, Zarinfar M, Rice S, Yang H, Yuan W, Camarero JA, Yu J, Chen XS, Zhang C, Feng P. 2021. SARS-CoV-2 Nsp5 Demonstrates Two Distinct Mechanisms Targeting RIG-I and MAVS To Evade the Innate Immune Response. mBio 12:e0233521.

51. Ye C, Chiem K, Park JG, Oladunni F, Platt RN, 2nd, Anderson T, Almazan F, de la Torre JC, Martinez-Sobrido L. 2020. Rescue of SARS-CoV-2 from a Single Bacterial Artificial Chromosome. mBio 11.

52. Tischer BK, Smith GA, Osterrieder N. 2010. En passant mutagenesis: a two step markerless red recombination system. Methods Mol Biol 634:421–430.

53. Chen DY, Khan N, Close BJ, Goel RK, Blum B, Tavares AH, Kenney D, Conway HL, Ewoldt JK, Chitalia VC, Crossland NA, Chen CS, Kotton DN, Baker SC, Fuchs SY, Connor JH, Douam F, Emili A, Saeed M. 2021. SARS-CoV-2 Disrupts Proximal Elements in the JAK-STAT Pathway. J Virol 95:e0086221.

54. Zhou R, To KK, Wong YC, Liu L, Zhou B, Li X, Huang H, Mo Y, Luk TY, Lau TT, Yeung P, Chan WM, Wu AK, Lung KC, Tsang OT, Leung WS, Hung IF, Yuen KY, Chen Z. 2020. Acute SARS-CoV-2 Infection Impairs Dendritic Cell and T Cell Responses. Immunity 53:864–877 e865.

55. Perez-Gomez A, Vitalle J, Gasca-Capote C, Gutierrez-Valencia A, Trujillo-Rodriguez M, Serna-Gallego A, Munoz-Muela E, Jimenez-Leon MLR, Rafii-El-Idrissi Benhnia M, Rivas-Jeremias I, Sotomayor C, Roca-Oporto C, Espinosa N, Infante-Dominguez C, Crespo-Rivas JC, Fernandez-Villar A, Perez-Gonzalez A, Lopez-Cortes LF, Poveda E, Ruiz-Mateos E, Virgen del Rocio Hospital C-WT. 2021. Dendritic cell deficiencies persist seven months after SARS-CoV-2 infection. Cell Mol Immunol 18:2128–2139.

56. Winheim E, Rinke L, Lutz K, Reischer A, Leutbecher A, Wolfram L, Rausch L, Kranich J, Wratil PR, Huber JE, Baumjohann D, Rothenfusser S, Schubert B, Hilgendorff A, Hellmuth JC, Scherer C, Muenchhoff M, von Bergwelt-Baildon M, Stark K, Straub T, Brocker T, Keppler OT, Subklewe M, Krug AB. 2021. Impaired function and delayed regeneration of dendritic cells in COVID-19. PLoS Pathog 17:e1009742.

57. Montoya M, Schiavoni G, Mattei F, Gresser I, Belardelli F, Borrow P, Tough DF. 2002. Type I interferons produced by dendritic cells promote their phenotypic and functional activation. Blood 99:3263–3271.

58. Skold AE, Mathan TSM, van Beek JJP, Florez-Grau G, van den Beukel MD, Sittig SP, Wimmers F, Bakdash G, Schreibelt G, de Vries IJM. 2018. Naturally produced type I IFNs enhance human myeloid dendritic cell maturation and IL-12p70 production and mediate elevated effector functions in innate and adaptive immune cells. Cancer Immunol Immunother 67:1425–1436.

59. Simmons DP, Wearsch PA, Canaday DH, Meyerson HJ, Liu YC, Wang Y, Boom WH, Harding CV. 2012. Type I IFN drives a distinctive dendritic cell maturation phenotype that allows continued class II MHC synthesis and antigen processing. J Immunol 188:3116–3126.

60. Schoggins JW. 2019. Interferon-Stimulated Genes: What Do They All Do? Annu Rev Virol 6:567–584.

61. Lee D, Le Pen J, Yatim A, Dong B, Aquino Y, Ogishi M, Pescarmona R, Talouarn E, Rinchai D, Zhang P, Perret M, Liu Z, Jordan I, Elmas Bozdemir S, Bayhan GI, Beaufils C, Bizien L, Bisiaux A, Lei W, Hasan M, Chen J, Gaughan C, Asthana A, Libri V, Luna JM, Jaffre F, Hoffmann HH, Michailidis E, Moreews M, Seeleuthner Y, Bilguvar K, Mane S, Flores C, Zhang Y, Arias AA, Bailey R, Schluter A, Milisavljevic B, Bigio B, Le Voyer T, Materna M, Gervais A, Moncada-Velez M, Pala F, Lazarov T, Levy R, Neehus AL, Rosain J, Peel J, Chan YH, Morin MP, Pino-Ramirez RM, Belkaya S, Lorenzo L, Anton J, Delafontaine S, Toubiana J, Bajolle F, Fumado V, DeDiego ML, Fidouh N, Rozenberg F, Perez-Tur J, Chen S, Evans T, Geissmann F, Lebon P, Weiss SR, Bonnet D, Duval X, Co VCCss, sign CHGEp, Pan-Hammarstrom Q, Planas AM, Meyts I, Haerynck F, Pujol A, Sancho-Shimizu V, Dalgard CL, Bustamante J, Puel A, Boisson-Dupuis S, Boisson B, Maniatis T, Zhang Q, Bastard P, Notarangelo L, Beziat V, Perez de Diego R, Rodriguez-Gallego C, Su HC, Lifton RP, Jouanguy E, Cobat A, Alsina L, Keles S, Haddad E, Abel L, Belot A, Quintana-Murci L, Rice CM, Silverman RH, Zhang SY, Casanova JL. 2023. Inborn errors of OAS-RNase L in SARS-CoV-2-related multisystem inflammatory syndrome in children. Science 379:eabo3627.

62. Stawowczyk M, Van Scoy S, Kumar KP, Reich NC. 2011. The interferon stimulated gene 54 promotes apoptosis. J Biol Chem 286:7257–7266.

63. Shi G, Kenney AD, Kudryashova E, Zani A, Zhang L, Lai KK, Hall-Stoodley L, Robinson RT, Kudryashov DS, Compton AA, Yount JS. 2021. Opposing activities of IFITM proteins in SARS-CoV-2 infection. EMBO J 40:e106501.

64. Prelli Bozzo C, Nchioua R, Volcic M, Koepke L, Kruger J, Schutz D, Heller S, Sturzel CM, Kmiec D, Conzelmann C, Muller J, Zech F, Braun E, Gross R, Wettstein L, Weil T, Weiss J, Diofano F, Rodriguez Alfonso AA, Wiese S, Sauter D, Munch J, Goffinet C, Catanese A, Schon M, Boeckers TM, Stenger S, Sato K, Just S, Kleger A, Sparrer KMJ, Kirchhoff F. 2021. IFITM proteins promote SARS-CoV-2 infection and are targets for virus inhibition in vitro. Nat Commun 12:4584.

65. Sposito B, Broggi A, Pandolfi L, Crotta S, Clementi N, Ferrarese R, Sisti S, Criscuolo E, Spreafico R, Long JM, Ambrosi A, Liu E, Frangipane V, Saracino L, Bozzini S, Marongiu L, Facchini FA, Bottazzi A, Fossali T, Colombo R, Clementi M, Tagliabue E, Chou J, Pontiroli AE, Meloni F, Wack A, Mancini N, Zanoni I. 2021. The interferon landscape along the respiratory tract impacts the severity of COVID-19. Cell 184:4953–4968 e4916.

66. Chang CH, Gourley TS, Sisk TJ. 2002. Function and regulation of class II transactivator in the immune system. Immunol Res 25:131–142.

67. Tsuchida T, Zou J, Saitoh T, Kumar H, Abe T, Matsuura Y, Kawai T, Akira S. 2010. The ubiquitin ligase TRIM56 regulates innate immune responses to intracellular double-stranded DNA. Immunity 33:765–776.

68. Zou L, Feng Y, Li Y, Zhang M, Chen C, Cai J, Gong Y, Wang L, Thurman JM, Wu X, Atkinson JP, Chao W. 2013. Complement factor B is the downstream effector of TLRs and plays an important role in a mouse model of severe sepsis. J Immunol 191:5625–5635.

69. Ben-Sasson SZ, Wang K, Cohen J, Paul WE. 2013. IL-1beta strikingly enhances antigen-driven CD4 and CD8 T-cell responses. Cold Spring Harb Symp Quant Biol 78:117–124.

70. Powell MD, Read KA, Sreekumar BK, Jones DM, Oestreich KJ. 2019. IL-12 signaling drives the differentiation and function of a T(H)1-derived T(FH1)-like cell population. Sci Rep 9:13991.

71. Bao L, Deng W, Huang B, Gao H, Liu J, Ren L, Wei Q, Yu P, Xu Y, Qi F, Qu Y, Li F, Lv Q, Wang W, Xue J, Gong S, Liu M, Wang G, Wang S, Song Z, Zhao L, Liu P, Zhao L, Ye F, Wang H, Zhou W, Zhu N, Zhen W, Yu H, Zhang X, Guo L, Chen L, Wang C, Wang Y, Wang X, Xiao Y, Sun Q, Liu H, Zhu F, Ma C, Yan L, Yang M, Han J, Xu W, Tan W, Peng X, Jin Q, Wu G, Qin C. 2020. The pathogenicity of SARS-CoV-2 in hACE2 transgenic mice. Nature 583:830–833.

72. Yinda CK, Port JR, Bushmaker T, Offei Owusu I, Purushotham JN, Avanzato VA, Fischer RJ, Schulz JE, Holbrook MG, Hebner MJ, Rosenke R, Thomas T, Marzi A, Best SM, de Wit E, Shaia C, van Doremalen N, Munster VJ. 2021. K18-hACE2 mice develop respiratory disease resembling severe COVID-19. PLoS Pathog 17:e1009195.

73. Anderson CK, Brossay L. 2016. The role of MHC class Ib-restricted T cells during infection. Immunogenetics 68:677–691.

74. Zhao Y, Cao Y, Chen Y, Wu L, Hang H, Jiang C, Zhou X. 2021. B2M gene expression shapes the immune landscape of lung adenocarcinoma and determines the response to immunotherapy. Immunology 164:507–523.

75. Vogl T, Stratis A, Wixler V, Voller T, Thurainayagam S, Jorch SK, Zenker S, Dreiling A, Chakraborty D, Frohling M, Paruzel P, Wehmeyer C, Hermann S, Papantonopoulou O, Geyer C, Loser K, Schafers M, Ludwig S, Stoll M, Leanderson T, Schultze JL, Konig S, Pap T, Roth J. 2018. Autoinhibitory regulation of S100A8/S100A9 alarmin activity locally restricts sterile inflammation. J Clin Invest 128:1852–1866.

76. Oh H, Ghosh S. 2013. NF-kappaB: roles and regulation in different CD4(+) T-cell subsets. Immunol Rev 252:41–51.

77. Suzuki AS, Yagi R, Kimura MY, Iwamura C, Shinoda K, Onodera A, Hirahara K, Tumes DJ, Koyama-Nasu R, Iismaa SE, Graham RM, Motohashi S, Nakayama T. 2020. Essential Role for CD30-Transglutaminase 2 Axis in Memory Th1 and Th17 Cell Generation. Front Immunol 11:1536.

78. Ricklin D, Reis ES, Mastellos DC, Gros P, Lambris JD. 2016. Complement component C3 - The "Swiss Army Knife" of innate immunity and host defense. Immunol Rev 274:33–58.

79. Jakos T, Pislar A, Jewett A, Kos J. 2019. Cysteine Cathepsins in Tumor-Associated Immune Cells. Front Immunol 10:2037.

80. Jian J, Konopka J, Liu C. 2013. Insights into the role of progranulin in immunity, infection, and inflammation. J Leukoc Biol 93:199–208.

81. Wang Q, Chen X, Feng J, Cao Y, Song Y, Wang H, Zhu C, Liu S, Zhu Y. 2013. Soluble interleukin-6 receptor-mediated innate immune response to DNA and RNA viruses. J Virol 87:11244–11254.

82. Yang XO, Pappu BP, Nurieva R, Akimzhanov A, Kang HS, Chung Y, Ma L, Shah B, Panopoulos AD, Schluns KS, Watowich SS, Tian Q, Jetten AM, Dong C. 2008. T helper 17 lineage differentiation is programmed by orphan nuclear receptors ROR alpha and ROR gamma. Immunity 28:29–39.

83. Miyazaki T, Muller U, Campbell KS. 1997. Normal development but differentially altered proliferative responses of lymphocytes in mice lacking CD81. EMBO J 16:4217–4225.

84. Stefanidakis M, Newton G, Lee WY, Parkos CA, Luscinskas FW. 2008. Endothelial CD47 interaction with SIRPgamma is required for human T-cell transendothelial migration under shear flow conditions in vitro. Blood 112:1280–1289.

85. Honda K, Yanai H, Negishi H, Asagiri M, Sato M, Mizutani T, Shimada N, Ohba Y, Takaoka A, Yoshida N, Taniguchi T. 2005. IRF-7 is the master regulator of type-I interferon-dependent immune responses. Nature 434:772–777.

86. Tannock LR, De Beer MC, Ji A, Shridas P, Noffsinger VP, den Hartigh L, Chait A, De Beer FC, Webb NR. 2018. Serum amyloid A3 is a high density lipoprotein-associated acute-phase protein. J Lipid Res 59:339–347.

87. Huntoon KM, Wang Y, Eppolito CA, Barbour KW, Berger FG, Shrikant PA, Baumann H. 2008. The acute phase protein haptoglobin regulates host immunity. J Leukoc Biol 84:170–181.

88. Aegerter H, Lambrecht BN, Jakubzick CV. 2022. Biology of lung macrophages in health and disease. Immunity 55:1564–1580.

89. Guo M, Morley MP, Jiang C, Wu Y, Li G, Du Y, Zhao S, Wagner A, Cakar AC, Kouril M, Jin K, Gaddis N, Kitzmiller JA, Stewart K, Basil MC, Lin SM, Ying Y, Babu A, Wikenheiser-Brokamp KA, Mun KS, Naren AP, Clair G, Adkins JN, Pryhuber GS, Misra RS, Aronow BJ, Tickle TL, Salomonis N, Sun X, Morrisey EE, Whitsett JA, Consortium NL, Xu Y. 2023. Guided construction of single cell reference for human and mouse lung. Nat Commun 14:4566.

90. Ivashkiv LB, Donlin LT. 2014. Regulation of type I interferon responses. Nat Rev Immunol 14:36–49.

91. Takeuchi O, Akira S. 2010. Pattern recognition receptors and inflammation. Cell 140:805–820.

92. Patil S, Fribourg M, Ge Y, Batish M, Tyagi S, Hayot F, Sealfon SC. 2015. Single-cell analysis shows that paracrine signaling by first responder cells shapes the interferon-beta response to viral infection. Sci Signal 8:ra16.

93. Jones CT, Catanese MT, Law LM, Khetani SR, Syder AJ, Ploss A, Oh TS, Schoggins JW, MacDonald MR, Bhatia SN, Rice CM. 2010. Real-time imaging of hepatitis C virus infection using a fluorescent cell-based reporter system. Nat Biotechnol 28:167–171.

94. Amor S, Fernandez Blanco L, Baker D. 2020. Innate immunity during SARS-CoV-2: evasion strategies and activation trigger hypoxia and vascular damage. Clin Exp Immunol 202:193–209.

95. Blanco-Melo D, Nilsson-Payant BE, Liu WC, Uhl S, Hoagland D, Moller R, Jordan TX, Oishi K, Panis M, Sachs D, Wang TT, Schwartz RE, Lim JK, Albrecht RA, tenOever BR. 2020. Imbalanced Host Response to SARS-CoV-2 Drives Development of COVID-19. Cell 181:1036–1045 e1039.

96. Alhammad YM, Parthasarathy S, Ghimire R, Kerr CM, O’Connor JJ, Pfannenstiel JJ, Chanda D, Miller CA, Baumlin N, Salathe M, Unckless RL, Zuniga S, Enjuanes L, More S, Channappanavar R, Fehr AR. 2023. SARS-CoV-2 Mac1 is required for IFN antagonism and efficient virus replication in cell culture and in mice. Proc Natl Acad Sci U S A 120:e2302083120.

97. Ferreira JC, Fadl S, Villanueva AJ, Rabeh WM. 2021. Catalytic Dyad Residues His41 and Cys145 Impact the Catalytic Activity and Overall Conformational Fold of the Main SARS-CoV-2 Protease 3-Chymotrypsin-Like Protease. Front Chem 9:692168.

98. Ricciardi S, Guarino AM, Giaquinto L, Polishchuk EV, Santoro M, Di Tullio G, Wilson C, Panariello F, Soares VC, Dias SSG, Santos JC, Souza TML, Fusco G, Viscardi M, Brandi S, Bozza PT, Polishchuk RS, Venditti R, De Matteis MA. 2022. The role of NSP6 in the biogenesis of the SARS-CoV-2 replication organelle. Nature 606:761–768.

99. Chang T, Yang J, Deng H, Chen D, Yang X, Tang ZH. 2022. Depletion and Dysfunction of Dendritic Cells: Understanding SARS-CoV-2 Infection. Front Immunol 13:843342.

100. Wang X, Guan F, Miller H, Byazrova MG, Cndotti F, Benlagha K, Camara NOS, Lei J, Filatov A, Liu C. 2023. The role of dendritic cells in COVID-19 infection. Emerg Microbes Infect 12:2195019.

101. Tahtinen S, Tong AJ, Himmels P, Oh J, Paler-Martinez A, Kim L, Wichner S, Oei Y, McCarron MJ, Freund EC, Amir ZA, de la Cruz CC, Haley B, Blanchette C, Schartner JM, Ye W, Yadav M, Sahin U, Delamarre L, Mellman I. 2022. IL-1 and IL-1ra are key regulators of the inflammatory response to RNA vaccines. Nat Immunol 23:532–542.

102. Chiem K, Ye C, Martinez-Sobrido L. 2020. Generation of Recombinant SARS-CoV-2 Using a Bacterial Artificial Chromosome. Curr Protoc Microbiol 59:e126.

103. Zagursky RJ, Hays JB. 1983. Expression of the phage lambda recombination genes exo and bet under lacPO control on a multi-copy plasmid. Gene 23:277–292.

104. Murphy KC. 1998. Use of bacteriophage lambda recombination functions to promote gene replacement in Escherichia coli. J Bacteriol 180:2063–2071.

105. Malherbe DC, Kurup D, Wirblich C, Ronk AJ, Mire C, Kuzmina N, Shaik N, Periasamy S, Hyde MA, Williams JM, Shi PY, Schnell MJ, Bukreyev A. 2021. A single dose of replication-competent VSV-vectored vaccine expressing SARS-CoV-2 S1 protects against virus replication in a hamster model of severe COVID-19. NPJ Vaccines 6:91.

106. Foley JW, Zhu C, Jolivet P, Zhu SX, Lu P, Meaney MJ, West RB. 2019. Gene expression profiling of single cells from archival tissue with laser-capture microdissection and Smart-3SEQ. Genome Res 29:1816–1825.

107. Ilinykh PA, Periasamy S, Huang K, Kuzmina NA, Ramanathan P, Meyer MN, Mire CE, Kuzmin IV, Bharaj P, Endsley JR, Chikina M, Sealfon SC, Widen SG, Endsley MA, Bukreyev A. 2022. A single intranasal dose of human parainfluenza virus type 3-vectored vaccine induces effective antibody and memory T cell response in the lungs and protects hamsters against SARS-CoV-2. NPJ Vaccines 7:47.

108. Meyer M, Wang Y, Edwards D, Smith GR, Rubenstein AB, Ramanathan P, Mire CE, Pietzsch C, Chen X, Ge Y, Cheng WS, Henry C, Woods A, Ma L, Stewart-Jones GB, Bock KW, Minai M, Nagata BM, Periasamy S, Shi PY, Graham BS, Moore IN, Ramos I, Troyanskaya OG, Zaslavsky E, Carfi A, Sealfon SC, Bukreyev A. 2021. Attenuated activation of pulmonary immune cells in mRNA-1273-vaccinated hamsters after SARS-CoV-2 infection. J Clin Invest 131.

109. Hao Y, Hao S, Andersen-Nissen E, Mauck WM, 3rd, Zheng S, Butler A, Lee MJ, Wilk AJ, Darby C, Zager M, Hoffman P, Stoeckius M, Papalexi E, Mimitou EP, Jain J, Srivastava A, Stuart T, Fleming LM, Yeung B, Rogers AJ, McElrath JM, Blish CA, Gottardo R, Smibert P, Satija R. 2021. Integrated analysis of multimodal single-cell data. Cell 184:3573–3587 e3529.

110. Bais AS, Kostka D. 2020. scds: computational annotation of doublets in single-cell RNA sequencing data. Bioinformatics 36:1150–1158.

111. Aran D, Looney AP, Liu L, Wu E, Fong V, Hsu A, Chak S, Naikawadi RP, Wolters PJ, Abate AR, Butte AJ, Bhattacharya M. 2019. Reference-based analysis of lung single-cell sequencing reveals a transitional profibrotic macrophage. Nat Immunol 20:163–172.

112. Kuleshov MV, Jones MR, Rouillard AD, Fernandez NF, Duan Q, Wang Z, Koplev S, Jenkins SL, Jagodnik KM, Lachmann A, McDermott MG, Monteiro CD, Gundersen GW, Ma’ayan A. 2016. Enrichr: a comprehensive gene set enrichment analysis web server 2016 update. Nucleic Acids Res 44:W90–97.

